# Allosteric capsid inhibitors and their escape mutants drive HIV-1 sensing

**DOI:** 10.64898/2026.08.21.746246

**Authors:** Kate L. Morling, Morten L. Govasli, Ben Graham, Justin Warne, Lauren Harrison, Lucy G. Thorne, Lydia S. Newton, Joshua Maw, Emma Touizer, Rebecca Sumner, Sally Oxenford, Joanna Rowley, Dara Annett, David Jacques, Till Böcking, Nikos Pinotsis, David L. Selwood, Greg J. Towers

**Affiliations:** Division of Infection and Immunity, University College London, London, UK; Wolfson Institute for Biomedical Research, University College London, London, UK; Cambridge Institute for Medical Research, University of Cambridge, Cambridge, UK; Department of Biomedicine, Centre for Cancer Biomarkers, University of Bergen, Bergen, Norway; The Institute of Cancer Research, Centre for Cancer Drug Discovery, Sutton, UK; Department of Infectious Disease, Sir Alexander Fleming Building, Imperial College Road, London, UK; Centre for Therapeutics Discovery, Lerner Research Institute, Cleveland Clinic, Cleveland, Ohio, US; Department of Infectious Diseases, King’s College London, Guy’s Hospital, London, UK; Centre for Immunobiology and Infection, Blizard Institute, Queen Mary University of London, Newark Street, London, UK; Department of Molecular Medicine, School of Biomedical Sciences, University of New South Wales, Sydney, New South Wales, Australia; Institute of Structural and Molecular Biology, School of Natural Sciences, Birkbeck College, London, UK

**Author notes:** These authors contributed equally (Kate L. Morling, Morten L. Govasli).

## Abstract

Small-molecule capsid inhibitors suppress HIV-1 infectivity by binding to capsid at the same site as FG motif-bearing host cofactors Sec24C, NUP153, CPSF6 and disordered nucleoporins residing in the nuclear pore complex central channel. We have used rational design to develop inhibitors called “allosteres” that target this pocket and inhibit HIV-1 infectivity. X-ray crystal structures of capsid/inhibitor complexes, reveal allosteric shifts upon inhibitor binding in the capsid C-terminal domain which impact the capsid lattice three-fold symmetry axis. Consistent with an uncoating mechanism, we find that allosteres cause HIV-1 to trigger innate immune response dependent on viral DNA and DNA sensor cGAS. Allosteres exhibit a similar loss of potency against clinically induced Lenacapavir resistance mutants but, strikingly, we find that HIV-1 bearing key resistance mutations induces innate immune activation in the absence of inhibitor. We hypothesise that resistant mutant sensitivity to cGAS contributes to reduction of HIV-1 transmission during Lenacapavir use in prophylaxis. Our work expands the physicochemical space and scaffold range for HIV capsid targeting inhibitors, provides mechanistic details of inhibition and facilitates improved inhibitor design.

## Introduction

HIV/AIDS continues to threaten public health with 1.3 million new infections and 630,000 deaths in 2024 (World Health Organization). In the absence of an effective vaccine or cure, prophylactic inhibitor regimens and treatment-as-prevention are expected to be the most effective routes to eradication. Thus, continued anti-HIV drug development is required. HIV is an exemplar virus that is extremely well characterised and understood, facilitating novel inhibitory approaches. For example, inhibiting host-virus protein-protein interactions (PPIs) should be an effective strategy because disturbing these interactions inhibits infection and can drive activation of natural innate immune defences (Sumner *et al*, 2020; Eschbach *et al*, 2024; Rasaiyaah *et al*, 2013).

The HIV capsid is central to infection and represents an under-exploited drug target. It is a molecular machine responsible for protecting and delivering the encapsidated viral genome to gene-rich host chromatin for integration. The capsid, or viral core, comprises around 250 hexamers and exactly 12 pentamers of the capsid protein (CA) (Perilla & Schulten, 2017). Here we refer to the capsid protein as CA and the viral core made of capsid protein as the capsid. Reverse transcription (RT) takes place inside the capsid, which shields newly synthesised viral DNA (vDNA) from innate immune detection by sensors including cyclic GMP-AMP synthase (cGAS) or degradation by nucleases including TREX1, facilitating innate immune evasion prior to uncoating in the nucleus for integration (Rasaiyaah *et al*, 2013; Sumner *et al*, 2020; Zuliani-Alvarez *et al*, 2022). Indeed, our earlier work suggested that a key feature of the single pandemic HIV-1 lineage, HIV-1(M), is specific adaptation of capsid to avoid TRIM5α and cGAS sensing (Zuliani-Alvarez *et al*, 2022).

HIV capsid recruits a series of cofactors, reviewed in (Morling *et al*, 2025), which we hypothesise act as molecular checkpoints, providing directionality and regulating the timing and location of capsid uncoating and genome release (Twarock *et al*, 2024). There are multiple distinct cofactor binding sites, with the first-identified being the loop on the capsid surface that recruits cofactor Cyclophilin A (CypA), which stabilises capsids in the cytoplasm (Ni *et al*, 2020; Briones *et al*, 2010), and NUP358, which assists nuclear transport and possibly viral trafficking (Schaller *et al*, 2011; Dharan *et al*, 2016). Separately, a hydrophobic pocket at the N-terminal domain (NTD)-C-terminal domain (CTD) interface between adjacent CA monomers recruits cofactors bearing phenylalanine-glycine (FG) motifs that hydrogen bond (H-bond) to CA N57 amongst other residues (Dickson *et al*, 2024; Rebensburg *et al*, 2021; Price *et al*, 2014) These include Sec24C in the cytoplasm, which stabilises capsids (Rebensburg *et al*, 2021), disordered nucleoporins, including NUP98, which permit capsids to phase-separate into the nuclear pore complex (NPC) for nuclear transport (Dickson *et al*, 2024; Fu *et al*, 2024), NUP153 in the inner nuclear pore (Price *et al*, 2014; Bhattacharya *et al*, 2014) and CPSF6 in the nucleus (Price *et al*, 2014), with evidence for CPSF6 recruiting capsids into phase-separated nuclear speckles to facilitate integration into transcriptionally active and gene-rich speckle-associated chromatin domains (SPADs) (Li *et al*, 2020; Luchsinger *et al*, 2023; Francis *et al*, 2020; Rensen *et al*, 2021; Achuthan *et al*, 2018; Schaller *et al*, 2011; Sowd *et al*, 2016). Disrupting these interactions, and thus breaking capsid stability, innate immune evasion, reverse transcription and nuclear import, is an attractive inhibitor strategy to inhibit HIV at multiple lifecycle stages.

The HIV-1 capsid FG binding site is druggable (Blair *et al*, 2010). A prototype capsid-targeting inhibitor called PF74, developed from a screen hit by Pfizer, has a phenyl group which mimics cofactor FG binding (Blair *et al*, 2010; Price *et al*, 2012, 2014; Bhattacharya *et al*, 2014; Gres *et al*, 2023) Several PF74-like analogues have been developed (Supplementary Fig 1) (Dostálková *et al*, 2020; Xu *et al*, 2018; Vernekar *et al*, 2020), with the first-in-class being Gilead Sciences’ Lenacapavir (LEN) (Yant *et al*, 2019; Link *et al*, 2020). LEN is approved by the FDA for adult people living with HIV (PLWH) with multidrug-resistant HIV. This heavily fluorinated molecule features a phenylalanine derived core decorated with an alkyne-sulphone arm, which contacts residues on the adjacent CA monomer. Critically, LEN is highly metabolically stable and effective with infrequent (6 month) injections (Link *et al*, 2020). Ongoing clinical trials also support LEN as a highly effective monotherapy prophylactic (Bekker *et al*, 2024).

Here we describe a series of independently-derived capsid targeting HIV inhibitors called allosteres. X-ray structures of inhibitor-CA complexes reveal an allosteric conformational change at the 2- and 3-fold symmetry axes in the capsid lattice, suggesting inhibitory mechanisms. Consistent with allosteres driving premature capsid uncoating in the cytoplasm, we find that allostere-treated infection drives innate immune activation dependent on viral DNA and sensor cGAS. Strikingly, untreated HIV bearing LEN/allostere resistance mutations activate cGAS sensing. We hypothesise that LEN prophylaxis benefits from LEN escape mutants being poor transmitters due to activation of innate immunity.

## Results

### Design of conformationally restricted capsid-targeting molecules that inhibit incoming HIV with nanomolar potency

We designed the allosteres to mimic binding of FG-containing cofactors to capsid. Pfizer’s PF74 has a tertiary amide which exists in solution as two isomers (*cis* and *trans*), with only the *cis* form binding CA (Fig 1A) (Bhattacharya *et al*, 2014). The phenylamide N-methyl group (orange) is expected to increase the proportion of the *cis* form (Manea *et al*, 1997), and we hypothesised its replacement with a cyclic 1,2,4-triazole (TRO-4, Fig 1B) would lock the inhibitor into the CA-binding conformation, improving potency, whilst retaining the phenyl core that mimics the FG cofactor interaction. The triazole core also provides conjugation points for additional CA binding groups. We therefore performed structure–activity relationship (SAR) studies measuring infectivity of VSV-G pseudotyped HIV-1 vector bearing a GFP-encoding genome (HIV-1 GFP) in U87 cells, measuring GFP positive cells by flow cytometry (Table 1, Fig 1C, Supplementary Fig 3A-C, Supplementary Table 1-3).

**Figure 1:**
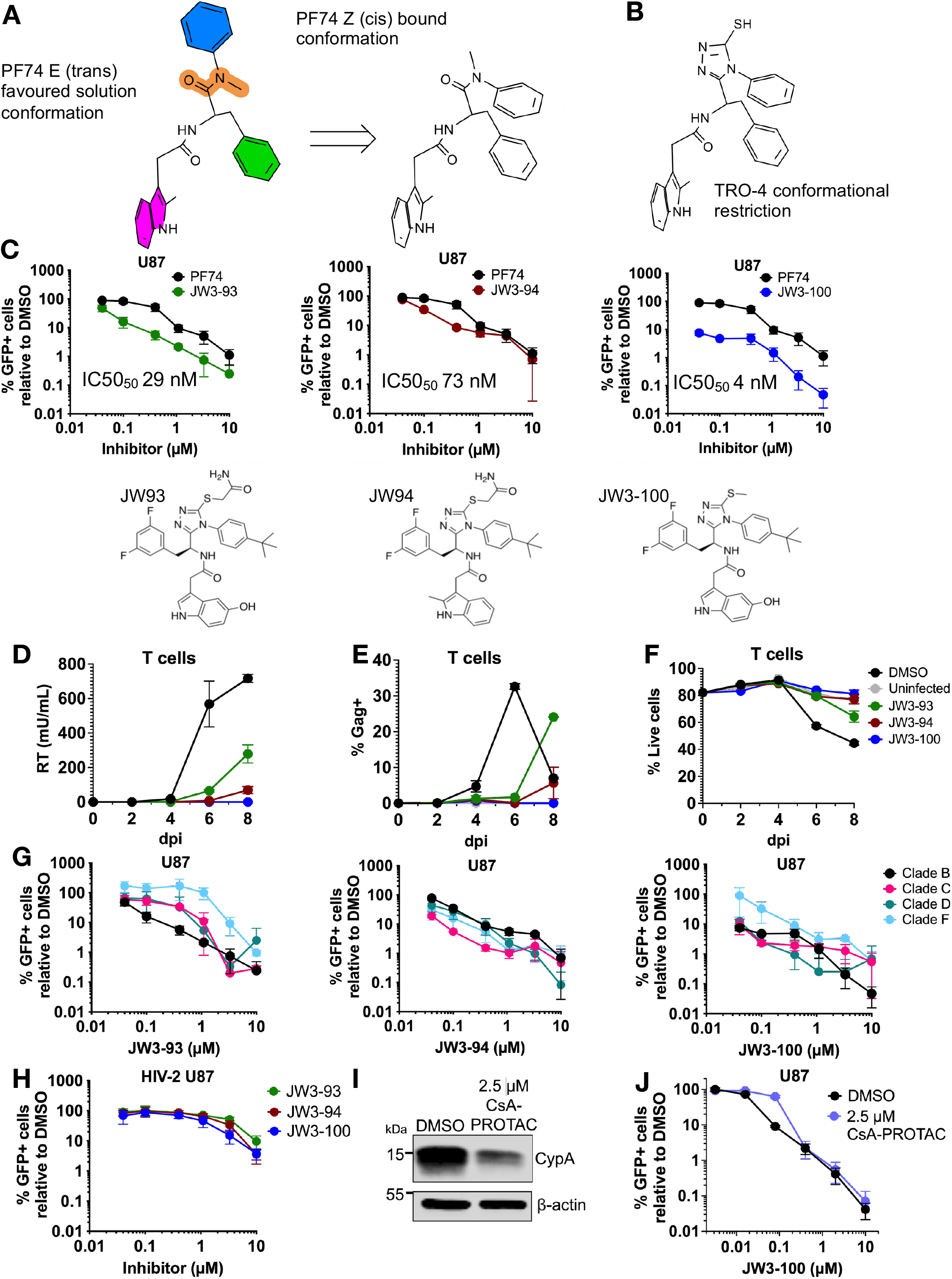
Conformationally restricted HIV-1 capsid inhibitors inhibit infection. **A** PF74 E (trans) and Z (cis) conformations. Chemical regions : indole group (pink), phenyl group (green), C-terminal cap (blue) and amide bond (orange). **B** Cyclised TRO-4 derivative with triazole ring. **C** U87 cells infected with HIV-1 GFP (MOI 0.25) after treatment with PF74 (black), JW3-93 (green), JW3-94 (red) or JW3-100 (blue) normalised to DMSO control value, mean ±SD (n=2). Inhibitor structures shown. **D-F** Primary CD4+ T-cells infected with HIV-1 NL4.3 were treated with 500 nM JW3-93/94/100, or DMSO, data at indicated days post infection, mean ± SD (duplicates for each donor), (**D**) Supernatant virus measured by SG-PERT, (**E**) % Gag+ cells (flow cytometry), (**F**) % Live cells (flow cytometry). Data from two more T-cell donors in Supplementary Figure 3. **G** % infected U87 cells measured as above after HIV-1 GFP infection (MOI 0.25) bearing Gag from clades B (R9, black), C (92BR025, pink), D (94UG114, teal) and F (93BR020, pale blue) and treatment with inhibitors as shown (n=2, mean ± SD). **H** % infected U87 cells, measured as above, after HIV-2 GFP infection (MOI 0.25) and treatment with inhibitors as shown, mean ± SD (n=2). I Immunoblot of U87 cells following 48 hours treatment with 2.5 μM CsA PROTAC JW4-10 or DMSO, detecting cyclophilin A (CypA) or β-actin as loading control (n=2, representative blot shown). **J** Corresponding proportion of GFP+ U87 cells, measured as above, after HIV-1 GFP infection (MOI 0.3) with JW3-100 following 48 hours pretreatment with 2.5 μM CsA PROTAC or DMSO, normalised to 0 μM JW3-100, mean ± SD (n=2 in technical triplicate).

**Table 1:** Lead allosteres from the SAR series. 12 allosteres with increased potency compared to PF74. Common structure with variable groups R_1_-R_5_ shown in table. An – 4-Anisole; t BuPh – 4-tert-Butylbenzene. KD determined by surface plasmon resonance (SPR) using crosslinked CA hexamers and calculated using kinetic and steady state affinity analysis. IC_50_ and IC_90_ determined by titration of allosteres on U87 cells during infection with VSV-G pseudotyped HIV1 GFP (MOI 0.25), measured by flow cytometry. Mean ± SD (n=3).

|  | R<sub>1</sub> | R<sub>2</sub> | R<sub>3</sub> | R<sub>4</sub> | R<sub>5</sub> | K<sub>D</sub> (SPR Kinetics) (nM) | K<sub>D</sub> (SPR Affinity) (nM) | IC<sub>50</sub> (μM) ±SD | IC<sub>90</sub> (μM) ±SD |
| --- | --- | --- | --- | --- | --- | --- | --- | --- | --- |
| **PF74** |  |  |  |  |  | 182 | 239 | 0.326 ± 0.042 | 1.236 ± 0.062 |
| **JW3-76** |  |  | Me | F | H | 65 | 867 | 0.192 ± 0.093 | 0.959 ± 0.171 |
| **JW3-79** | An | Me | H | F | OH | 468 | 811 | 0.073 ± 0.071 | 0.289 ± 0.102 |
| **JW3-82** | An | CH<sub>2</sub>CONH<sub>2</sub> | Me | F | H | 109 | 995 | 0.292 ± 0.129 | 1.021 ± 0.056 |
| **JW3-93** | <sup>t</sup>BuPh | CH<sub>2</sub>CONH<sub>2</sub> | H | F | OH | 62 | 2849 | 0.029 ± 0.057 | 0.151 ± 0.088 |
| **JW3-94** | <sup>t</sup>BuPh | CH<sub>2</sub>CONH<sub>2</sub> | Me | F | H | 133 | 2275 | 0.073 ± 0.060 | 0.257 ± 0.061 |
| **JW3-100** | <sup>t</sup>BuPh | Me | H | F | OH | 256 | 1261 | 0.004 ± 0.072 | 0.020 ± 0.164 |
| **JW3-132** | <sup>t</sup>BuPh | CH<sub>2</sub>CONHMe | H | F | OH | 58 | 3821 | 0.042 ± 0.063 | 0.541 ± 0.125 |
| **JW3-133** | <sup>t</sup>BuPh | CH<sub>2</sub>CONHMe | Me | F | H | 202 | 2902 | 0.115 ± 0.056 | 0.596 ± 0.096 |
| **JW3-134** | <sup>t</sup>BuPh | CH<sub>2</sub>CONHMe | Me | F | F | 416 | 3553 | 0.053 ± 0.068 | 0.157 ± 0.048 |
| **JW3-135** | <sup>t</sup>BuPh | CH<sub>2</sub>CONMe<sub>2</sub> | H | F | OH | 66 | 4844 | 0.050 ± 0.058 | 0.217 ± 0.071 |
| **JW3-136** | <sup>t</sup>BuPh | CH<sub>2</sub>CONH<sub>2</sub> | Me | F | H | 133 | 2275 | 0.073 ± 0.060 | 0.257 ± 0.061 |
| **JW3-137** | <sup>t</sup>BuPh | CH<sub>2</sub>CONMe<sub>2</sub> | Me | F | F | 804 | 4418 | 0.067 ± 0.111 | 0.303 ± 0.055 |

TRO-4 inhibited infection (Supplementary Table 1), and further SAR candidates were prepared through an optimised linear synthesis route (Supplementary Fig 2). Briefly, SAR established that phenyl C-terminal caps at R1 were advantageous and thiols/thioethers were tolerated at R2 (Supplementary Table 1). Fluorination has been shown to improve the activity of PF74 derivatives (Jiang *et al*, 2019) By varying the C-capping group at R1 from phenyl to anisole and with fluorination of the benzyl group at R4, we increased binding and antiviral potency (Supplementary Table 3). Replacement of R1 with a para-*^t^*Butyl benzene motif led to the discovery of a series of highly potent analogues, 12 of which had greater potency than PF74 (Table 1, Supplementary Table 3). JW3-100, which possesses a methyl thioether side chain and a 3-hydroxyindole motif, emerged as our most potent inhibitor (IC_50_ 4 nM). JW3-93 (IC_50_ 29 nM) and JW3-94 (IC_50_ 73 nM) were also selected for further characterisation (Fig 1C).

Intriguingly, allosteres exhibited different shaped inhibition curves. JW3-100 displayed a biphasic curve, whilst JW3-93 was monophasic, and JW3-94 was triphasic suggesting subtle differences between inhibitory mechanisms despite similar potencies (Fig 1C). All three inhibitors suppressed HIV-1 NL4-3 replication in primary CD4+ T-cells with minimal toxicity (Fig 1D-F, Supplementary Fig 3D-I). Surprisingly, whilst JW3-93 was more potent than JW3-94 in single-round infection in U87 cells, JW3-94 more effectively inhibited HIV-1 spreading infection in T-cells, suggesting different modes of inhibition can underlie potency differences across cell types and infection models. Allosteres displayed activity against HIV-1 GFP bearing Gag from multiple HIV-1 subtypes (Zuliani-Alvarez *et al*, 2022; Ikeda *et al*, 2004; Gao *et al*, 1998) (C, D and to a lesser extent F) (Fig 1G, Supplementary Table 4), consistent with only 8% CA FG pocket variation between subtypes (Supplementary Fig 4). We found little activity against HIV-2, consistent with SIVsm/HIV-2 lineage viruses’ divergent CA sequences and cofactor dependencies (Fig 1H) (Schaller *et al*, 2011; Mamede *et al*, 2017)

CypA interacts with a different site on CA to allosteres but there is evidence for allosteric interactions between the CypA and FG capsid binding sites (Zuliani-Alvarez *et al*, 2022; Twarock *et al*, 2024; Lu *et al*, 2015). Concordantly, we found that if we degraded CypA using a cyclosporine A proteolysis targeting chimera (CsA-PROTAC) (Colpitts *et al*, 2020) (Fig 1I), JW3-100 became less effective against HIV-1 GFP at low concentrations, shifting the JW3-100 IC_50_ from 27 nM to 93 nM (Fig 1J). Therefore, CypA appears to be important for allostere antiviral activity, with CypA loss shifting the inhibition curve shape from biphasic to monophasic. This observation reflects similar CypA sensitivity for CA inhibitor PF74 (Shi *et al*, 2011).

### Allosteres interfere with production of infectious HIV virions

CA inhibitors PF74 and LEN interfere with HIV-1 particle production (Yant *et al*, 2019; Link *et al*, 2020; Blair *et al*, 2010; Huang *et al*, 2024) Similarly, allosteres impacted the infectivity of HIV-1 pLAI ΔEnv GFP produced in HEK293T cells. Measurement of viral particles released from cells treated with JW3-93/100 by reverse transcriptase assay (SG-PERT) indicated a small (2-fold) reduction in reverse transcriptase activity (Fig 2A). Concordantly, measurement of Gag expression (immunoblot) in the treated producer cells suggested that Gag expression was also not particularly allostere-sensitive (Fig 2B). Observed reduction of cleaved CA levels in cell extracts likely reflects decreased producer cell re-infection because Gag cleavage occurs after viral particle release (Sundquist & Krausslich, 2012). Measurement of Gag/CA in viral supernatants suggested only slightly reduced levels of CA (Fig 2C). However, JW3-93/100 treatment during production reduced the infectivity of HIV-1 GFP in U87 cells by over 50-fold (Fig 2D). In this experiment, virus preps were diluted by 1000x to ensure the residual inhibitor did not directly impact incoming infection and left for at least 15 minutes to allow for allostere dissociation (Supplementary Fig 11F). These data suggest allosteres do not particularly reduce Gag expression or the number of particles produced, but JW3-93 and JW3-100 strongly reduce the infectivity of viral particles produced in their presence. One possible mechanism for this is that allostere stabilise hexamers but destabilise pentamers, resulting in malformed and less infectious particles. Indeed, allosteres are expected to interact more weakly (or not at all) with pentamers because the binding site contains an additional 3_10_ helical turn, restructuring the pocket (Stacey *et al*, 2023; Schirra *et al*, 2023) Furthermore, LEN impairs pentamer formation whilst inducing assembly of hexameric lattices (tubes), explaining its inhibition of HIV-1 maturation (Huang *et al*, 2024).

**Figure 2:**
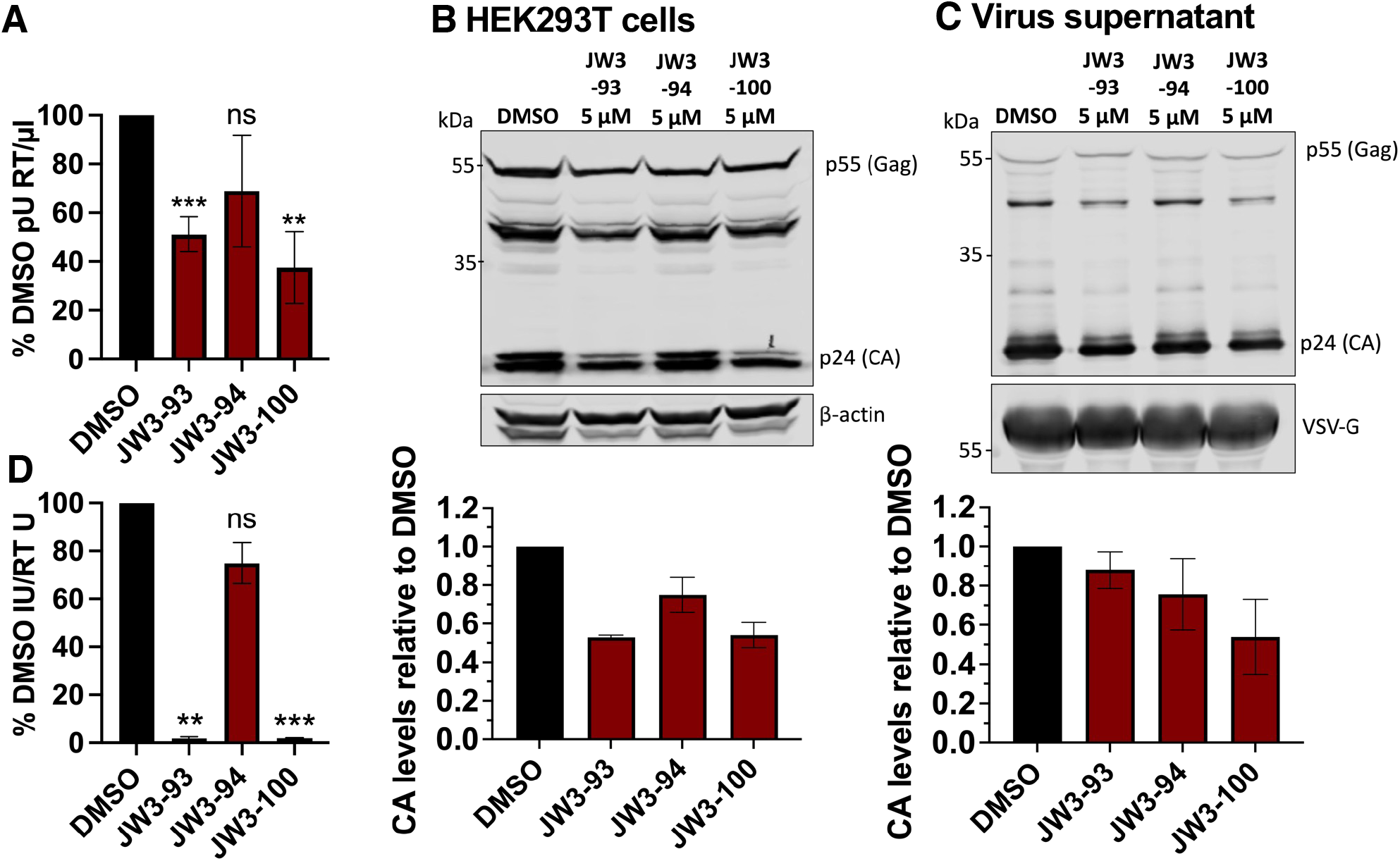
Allosteres interfere with production of infectious HIV-1. **A-D** HIV-1 LAI GFP produced in 293T cells treated with 5 μM allostere or DMSO vehicle and **A** supernatant reverse transcriptase (RT) activity (SG-PERT). Welch’s t-test, mean ± SD (n=4). **B** Immunoblot of 293T extracts detecting p24 or actin as loading control. CA protein quantification was normalised to actin and to DMSO values. mean ± SD (n=2). **C** Immunoblot of 293T supernatants, detecting p24 or VSV-G as loading control. Quantification of CA protein levels was normalised to VSV-G and to DMSO values mean ± SD (n=2). **D** % GFP+ U87 cells determined (flow cytometry) 48 hours post infection after a 1000-fold dilution with infectious units (IU) per ml normalised to RT activity. Welch’s t-test, ± SD (n=2 infections in technical triplicate).

### Structural comparison of HIV-1 CA in complex with allosteres and cofactor peptides reveals allosteric changes mediated via CA lysine 182

To better understand inhibitor-induced allosteric changes and antiviral mechanism, we determined X-ray crystal structures of native (not cross-linked) HIV-1 CA hexamers (1.97 Å) and hexamers in complex with allosteres JW3-76 (2.09 Å), JW3-93 (2.60 Å), JW3-94 (2.36 Å), JW3-100 (2.07 Å), and JW3-134 (2.50 Å) (Fig 3A-D and Supplementary Fig 6A-B). In addition, we solved structures of native CA hexamers in complex with peptides from CPSF6_Pro313-Gly327_ (2.40 Å) and NUP153_Thr1407-Thr1423_ (2.60 Å) which enabled us to directly compare allostere-induced conformational changes in CA with those caused by FG cofactor recruitment under the same experimental setup (Fig 3G, Supplementary Figs 5-8, Supplementary Table 6). We found that allosteres bind to the CA in a similar configuration, with their active groups mimicking the native CA/PF74 complex (PDB ID 4XFZ) (Gres *et al*, 2023, 2015). Each inhibitor is designed so that three of its functional groups - difluorophenyl, triazol or propenamide, and t-butyl-phenyl or methoxyphenyl form distinct interactions with the CA NTD thereby contributing to the binding affinity to a single CA molecule. Critically, the indole groups interact with both the CA NTD and the CTD of the adjacent CA’, likely interfering with the capsid lattice structure and explaining inhibition of infection (Fig 3A-E, Supplementary Fig 6A-B). However, the capsid– allostere complex structures are highly similar to one another (RMSD 0.32–0.51 Å) and to the unbound apo CA structure (RMSD 0.60–0.65 Å). Peptide-bound conformations show slightly greater divergence than inhibitor-bound states (RMSD 0.578-0.943 Å), (Supplementary Table 7), but differences are local and subtle.

**Figure 3:**
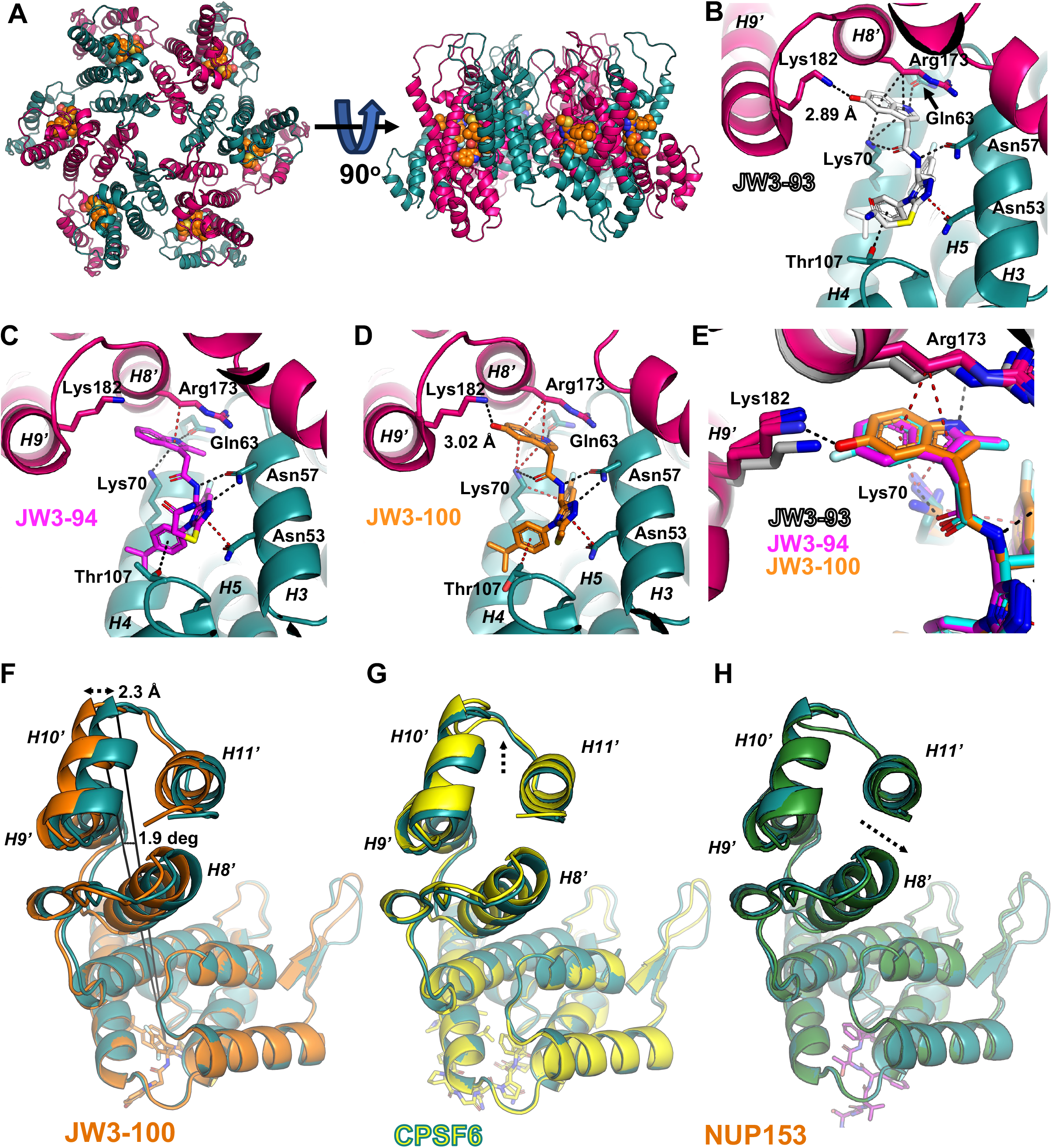
HIV1-CA hexamers with Allosteres and CPSF6. **A** Top and side view of the CA hexamer. Capsid protomers alternate pink/teal. Bound JW3-100 is shown as orange spheres. **B,C,D** Cartoon/stick representation of allostere binding interfaces JW3-93 (white), JW3-94 (magenta), and JW3-100 (orange). The main interacting residues and the helices (coloured as in panel A) are indicated. **E** Comparison of the three allostere binding motifs (JW3-93, JW3-94, JW3-100) to the native CA hexamer (in grey). The interacting Lys182 from the adjacent hexamer is coloured as corresponding inhibitor. Black dashed lines: H-bonds/electrostatic interactions. Red dashed lines: π-CH/ π-ion interactions. Summary Cartoon of binding effects as described in text. **F** Conformational changes in the CACTD α-helices H8’–H11’ in the JW3-100–bound structure (orange) compared with native ligand-free CA (green). The tilt angle and the maximum deviation at the distal end between the native and the allostere bound structure are shown **G** Conformational changes in the CACTD α-helices H8’–H11’ in the CPSF6–bound structure (yellow) compared with native ligand-free CA (green). The dashed arrow indicates a minimum shift of about 0.5 Å at the C-terminal helices of the CPSF6-bound structure compared to the native. **H** Conformational changes in the CACTD α-helices H8’–H11’ in the NUP153–bound structure (purple) compared with native ligand-free CA (green). The dashed arrow indicates the shift direction at the C-terminal helices of the NUP153-bound structure compared to the native.

Allosteres are designed to improve inhibitor affinity for the FG binding pocket on CA hexamers whilst mimicking cofactor binding, in particular, to the potential uncoating regulator CPSF6. The fluorinated phenyl groups replace the phenylalanine of the cofactor FG motif, strengthening hydrophobic contacts with Leu56, Met66, Leu69 and Ile73 and introducing a new π–CH interaction with Lys70 (Supplementary Fig 6C). Incorporation of a triazole ring locks inhibitors in the Z conformation, shortens the H-bond with Asn57, and adds a π–CH contact with Asn53 (Supplementary Fig 6). At the C-terminal cap, trimethyl (JW3-93/94/100/134) or methoxy (JW3-76) substitutions slightly shift positioning of the phenyl ring but preserve the Thr107 π–CH contact while the trimethyl group engages with Asn74 electrostatically (Fig 3B-D, Supplementary Fig 6A-D). At the same position, LEN forms two H-bonds with Asn74, and its methylsulfonyl group forms an extra H-bond with Thr54 consistent with a much tighter LEN-CA binding compared to allosteres (PDB ID 6V2F) (Link *et al*, 2020) (Supplementary Fig 6E, F).The -SR2 groups on the triazole ring may have a slight effect on the binding comparing similar allosteres - larger -SR2 groups such as in JW3-93 interact more closely with the loop between H5 and H6 while shorter groups such as in JW3-100 shift the entire inhibitor away from this loop and closer to the adjacent CA’.

A key observation may explain allosteric inhibitory activity. Inhibitor interaction with Lys182 of the adjacent CA’ induces a minor shift of the H9’ α-helix C-terminus, which propagates to the entire CTD of CA’, affecting the two- and three-fold inter-hexameric contacts that are key to viral core integrity (Zhao *et al*, 2013) (Fig 3B-E and Supplementary Fig 6A,B). Overlaying the structures with the CA FG binding sites aligned (Fig 3F-H) reveals that displacement of Lys182 (Fig 3E) alters the positioning of helix H9’ thereby perturbing the two-fold inter-hexamer interface (Supplementary Fig 5B). The resulting shift in H9’ propagates to H10’ and H11’ where these two C-terminal α-helices form the three-fold interface (Fig 3F, Supplementary Fig 8A, D). Thus, a local perturbation at Lys182 near the N-terminus of H9’ triggers a cascade of progressively larger conformational changes of up to about 2 Å (depending on the bound allostere) and 2 degrees tilt, constituting the observed allosteric effect (Fig 3F). Allosteres interact with Lys182 via two distinct modes. Those carrying a hydroxyl or fluorine group on the benzene ring of the indole core (JW3-93, JW3-100 and JW3-134) form a relatively strong H-bond with the amino group of Lys182. In contrast, methyl-substituted indoles on pyrrole ring (JW3-76, JW3-94, PF74 and partially JW3-134) shift the indole closer to CA’, creating steric accommodation within the binding pocket and resulting in hydrophobic repulsion with Lys182 (H-bond for JW3-134) (Fig 3B-E and Supplementary Fig 6A-B).

Beyond Lys182, allosteres engage three additional CA residues to influence CA-CA interactions. A H-bond (2.7–3.1 Å) is observed between the carboxyl group of the Gln63 side chain and the NH of the indole’s pyrrole ring in all allosteres (2.8 Å for JW3-100) (Fig 3B-C and Supplementary Fig 6A-B). In addition, Lys70 from the CA and Arg173 from the adjacent CA’ effectively “sandwich” the allostere indole, forming π–CH or π–electrostatic interactions with its aromatic rings (Fig 3B-E). Notably, only the hydroxyl-indole allosteres JW3-93 and JW3-100 interact with both their indole aromatic rings (Fig 3B-E). In contrast, the methyl-indole allosteres (JW3-76, JW3-94) interact like PF74, solely with the pyrrole ring, while the methyl-fluoro indole allostere (JW3-134) engages only the benzene ring (Fig 3B-E and Supplementary Fig 6A-B). Combined, these structures suggest that substitutions on the indole ring modulate its position, thereby influencing interactions with key CA and CA’ residues. The hydroxyl-indole substituent appears to promote a more favourable binding mode - potentially through enhanced engagement of the hydroxyl-indole group with both CA’ residues Lys182 and Arg173 - consistent with the higher potency observed for JW3-100 and JW3-93 over JW3-94, particularly regarding impact on particle infectivity (Fig 2D).

The crystal structures of CA in complex with CPSF6 and NUP153 peptides reiterate previously described key conformational changes in the hexagonal lattice induced by ligand binding (Supplementary Figs 7 and 8) (Gres *et al*, 2023; Bhattacharya *et al*, 2014; Price *et al*, 2014; Wei *et al*, 2022; Schirra *et al*, 2023; Stacey *et al*, 2023). The CPSF6 peptide utilizes an FG-binding motif (Pro313-Gly327) and CA Lys70 as the key residues for this interaction, while a four-residue interface (CPSF6 Phe316-Gln319) interacts with the adjacent CA’ α-helices H8’ and H9’. The interaction with H9’ is primarily electrostatic with the CPSF6 residues Gln319, Gly318 and Phe316 forming H-bonds and π-anion interactions with the side chains of Gln179, Lys182 and Asn183 of H9’ (Supplementary Fig 7A-B). Notably, engagement of Lys182 by CPSF6 mimics allosteric inhibitors shifting CA′ H8′ and H9′ α-helices to a smaller degree than the inhibitors, and unlike the allosteres, CPSF6 does not move the C-terminal H10’ and H11’ (Fig 3G and Supplementary Fig 8A-B) (Supplementary text). In contrast, the NUP153 peptide exerts a markedly different effect on CA, despite binding to the same pocket. Although the peptide used for crystallisation encompassed NUP153 residues 1407-1423, electron density is observed only for residues 1409-1418, with the last clearly resolved residues being Phe1417-Gly1418, which bind within the FG-binding pocket of CA. The preceding residues (1409-1416) extend through the cleft between the NTD and CTD of the adjacent CA’ toward the FG-binding pocket, forming only a single H-bond between NUP153 Ser1412 and the main chain carbonyl group of CA’ Gln176, located in the loop connecting H8’ and H9’ (Supplementary Fig 7C-D). This limited interaction suggests a relatively weak capsid-NUP153 association, consistent with the requirement for NUP153 to release the capsid following transit through the nuclear pore after infection. Notably, and unlike allosteres and CPSF6, NUP153 does not interact with CA Lys182 but causes a minor displacement of the CTD relative to the NTD of CA’ (Fig 3H and Supplementary Fig 8B-C).

Further analyses of CA allostery reveal that the small-molecule inhibitors induce a uniform hinge-like closure of the CA NTD and CTD, resulting in subtle displacements of CTD helices H8 and H9 involved in the two-fold inter-hexamer interface, and more pronounced displacements at the distal CTD helices H10 and H11 that form the three-fold inter-hexamer interface (Fig 3D, F, Supplementary Fig 8A & C, Supplementary Table 10). The host-factor peptides elicit smaller, more complex, motions: CPSF6 produces localized rearrangements mainly at helices H8 and H9 forming the two-fold interface and minor rearrangements at helices H10 and H11 (Fig 3G and Supplementary Fig 8B), while NUP153 induces a subtle but uniform shift of the entire CTD, resulting in a higher overall RMSD despite the limited magnitude of the local structural changes (Fig 3H & Supplementary Fig 8C middle panel). Thus, despite sharing a conserved FG-binding mode, allosteres and host factors differentially remodel inter-hexamer interfaces, providing a structural basis for their distinct effects on capsid stability and infectivity (see supplemental text for further details). Strikingly, LEN has been described to induce a similar hinge motion to allosteres, even with a crosslinked CA (Bester *et al*, 2020) and PF74 also induces similar allosteric shifts in the CA CTD relative to the NTD. (Supplementary Fig 8C and Supplementary Table 10).

To further assess the impact of ligand binding on the CA lattice, we examined the effect on the two- and three-fold symmetry axes of the inter-hexameric lattice in the crystals, which mimic the major hexamer interfaces in the viral capsid. Although crystal packing restricts large-scale movements through inter-plane contacts, the principal interaction outside the hexameric plane involves flexible loops connecting the CTD and NTD of CAs in the different layers. Thus, conformational changes observed within the crystal lattice are expected to approximate those occurring in the native capsid lattice. Analysis using the EBI PDBePISA server (Krissinel & Henrick, 2007), illustrates that the two-fold interface is consistently robust across all structures, featuring large surface areas and highly negative enthalpy - ΔG values. Notably, NUP153 promotes a more specific interaction, exhibiting by far the lowest P-value (0.066) and the most negative ΔG (−9.8 kcal/mol) (Supplementary Table 8) indicating that NUP153 significantly stabilizes the two-fold interface. Strikingly, enhanced stability is achieved without H-bonding, suggesting that hydrophobic interactions and/or shape complementarity, rather than electrostatics, are the dominant stabilizing forces. Conversely, CPSF6 yields the smallest two-fold interface area and a ΔG identical to the apo capsid. By contrast, allosteres partially mimic host factor binding but do not stabilize the two-fold interface as strongly as NUP153.

More detailed analysis of the two-fold interface reveals that NUP153 increases the enthalpic contribution of almost every amino acid involved in the interface. However, two distinct amino acid categories emerge: (i) The hydrophobic anchor “base” of the interface comprising Leu151, Val181, and Trp184, which provides a large and consistent contribution across all structures (values >1.0). (ii) The determinants of ligand specificity comprise Ser178 and Glu180, with Glu180 emerging as the most critical “switch” (Supplementary Table 9). In particular the enthalpic gain for NUP153 is the strongest; JW3-76 closely mimics this interaction, whereas JW3-100 fails to reproduce it altogether (Supplementary Table 8). At the three-fold interface, only CPSF6 transforms the contact into a statistically specific and structurally significant interaction, by more than doubling the interface area and enthalpic gain relative to the apo capsid. This enhancement is driven primarily by contributions from Ala204 and Leu205 (Supplementary Table 9). In contrast, NUP153 shows the opposite effect, inducing the smallest three-fold interface area, with minimal contribution from those residues, which is significantly lower than the unbound apo capsid. Notably, allosteric inhibitors reverse the directionality of the three-fold interactions, consistent with the uniform hinge-like closure they induce (Supplementary Fig 8A,C).

Within the context of the hexagonal lattice, CPSF6 may act as quasi-stabilising factor at the vertices where hexamers converge, effectively promoting local lattice rigidity through a subtle repositioning of the two C-terminal α-helices (Fig 3G, Supplementary Fig 8D). Biologically, CPSF6 has been proposed to regulate the uncoating process, because its manipulation alters integration targeting and innate immune sensing of infection (Schaller *et al*, 2011; Rasaiyaah *et al*, 2013). These data suggests that it does so, not just by binding, but by physically influencing the three-fold vertices of the fullerene cone. NUP153, by contrast, interacts with the capsid at the inner nuclear pore, likely during the translocation process. A minimized or “loosened” three-fold vertex might permit the capsid to deform as it traverses the pore. Allosteres fail to reinforce the capsid lattice at three-fold interfaces, as reflected by comparable interfacial areas and ΔG values to the apo capsid. However, their effect appears to arise from stabilizing an alternative CTD conformation that constrains the C-terminal helices in different positions from the apo structure (Supplementary Fig 8 A,C,D).

Overall, our data suggest that host factors modulate HIV capsid stability through distinct and spatially differentiated mechanisms: NUP153 rigidifies the individual hexamer building blocks at the two-fold interface, whereas CPSF6 locks the global fullerene lattice together by reinforcing the three-fold vertices. In contrast, allosteric inhibitors occupy the same binding pocket but fail to induce either of these infection-promoting effects, allowing them to act as competitive inhibitors of cofactor binding whilst likely having additional negative impact on regulation of capsid stability (Supplementary Fig 8D).

### Allosteres cause innate immune sensing of HIV-1 by cGAS

Previous work on PF74 and LEN, together with the presented structures of allosteres bound to CA hexamers, suggest allosteres may cause HIV capsid uncoating. Indeed, both PF74 and LEN drive premature capsid uncoating and genome release because the stabilised hexamers are flattened disturbing curvature and therefore core integrity (Faysal *et al*, 2024; Márquez *et al*, 2018; Selyutina *et al*, 2022; Christensen *et al*, 2020; Shi *et al*, 2011; Blair *et al*, 2010; Bhattacharya *et al*, 2014). In addition, they mimic/interfere with binding of host cofactors that regulate capsid stability. This could cause innate immune activation through premature release of viral genomes and sensing by cGAS. Certainly, both PF74 and LEN drive cGAS-dependent innate immune activation (Sumner *et al*, 2020; Kumar *et al*, 2018; Eschbach *et al*, 2024). As intact capsids are required for effective DNA synthesis in cells (Christensen *et al*, 2020; Jennings *et al*, 2020; Sowd *et al*, 2021), allosteres may also be expected to inhibit RT by driving uncoating.

To probe this, we used VSV-G pseudotyped HIV-1 ΔEnv with GFP in place of Nef (HIV-1 LAI GFP), and THP-1 cells which can sense VSV-G pseudotyped non-pandemic HIV and PF74-treated HIV effectively (Zuliani-Alvarez *et al*, 2022; Sumner *et al*, 2020). We used THP-1 cells bearing a luciferase reporter controlled by the endogenous IFIT1 promoter (Mankan *et al*, 2014) with a fixed dose of DNase-treated, sucrose-purified HIV-1 LAI GFP, in the presence of allosteres (0.1-10 μM JW3-93/94/100). We measured infected cells by flow cytometry (Fig 4A), induction of the IFIT1 reporter (luciferase, Fig 4B and supplementary Fig 9A), induction of ISG mRNAs (MxA, IFIT2 and CXCL10, qRT-PCR, Fig 4C and supplementary Fig 9B), and viral DNA synthesis (qPCR, Fig 4D and supplementary Fig 9C). In these experiments, we used Nevirapine (RT inhibitor) and Raltegravir (integrase inhibitor) as positive controls and boiled virus as a negative control for viral DNA synthesis. Whilst HIV-1 LAI GFP alone caused some ISG induction, this was enhanced by allostere treatment (Fig 4B and C and supplementary Fig 9A and B), supporting our hypothesis that allosteres break capsid open at the wrong time and place, activating innate immune responses. Allosteres caused activation to different extents, with JW3-94 causing the greatest innate immune activation at 1 μM and JW3-93 causing the least, likely influenced by the amount of viral DNA pathogen-associated molecular pattern (PAMP) present. Indeed, inhibition of DNA synthesis by allosteres was inversely correlated with sensing, with JW3-94 inhibiting DNA synthesis the least and JW3-93 inhibiting DNA synthesis the most (Fig 4D and supplementary Fig 9C). Similarly, with JW3-94 and JW3-100, peak innate immune activation was seen at an intermediate dose (1 μM), where we see sub-optimal inhibition of DNA synthesis, consistent with capsids being broken open after enough DNA has been synthesised to facilitate cGAS sensing at this dose. All three allosteres completely inhibited infection at 10 μM, where we observed little innate immune activation, suggesting high dose inhibitor breaks capsids open prior to DNA synthesis thereby preventing viral DNA formation and its sensing. Concordantly, no RT products were observed at this dose. All three allosteres similarly reduced 2-LTR circle formation (Fig 4D), which reflects inhibited viral nuclear transport, consistent with inhibition of nuclear pore associated FG bearing proteins binding CA (Fu *et al*, 2024; Dickson *et al*, 2024).

**Figure 4:**
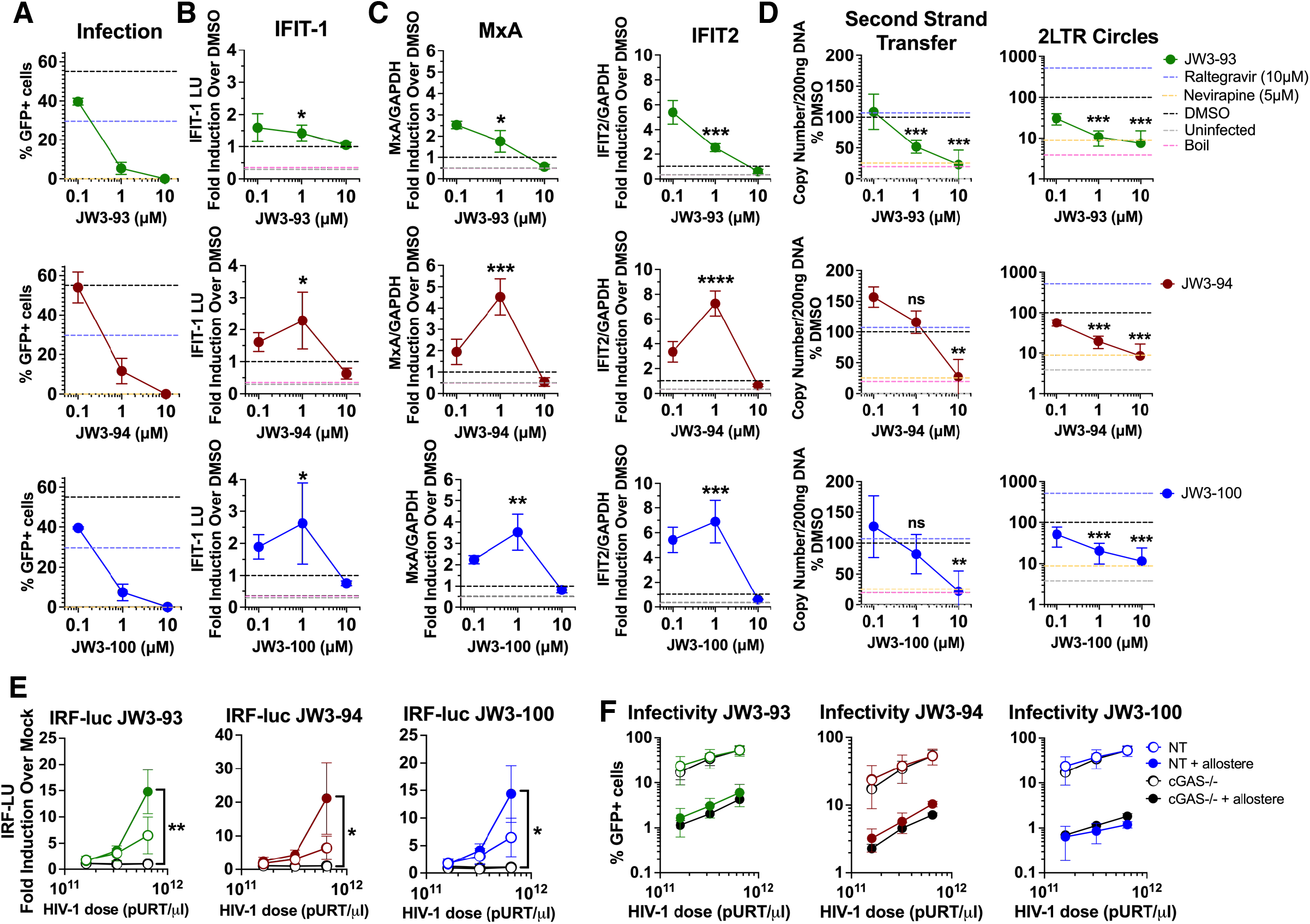
Allosteres cause innate immune sensing of HIV-1. **A** GFP+ THP-1 reporter cells measured 48 hours after HIV-1 LAI GFP (MOI 0.4) infection and treatment with JW3-93 (green), JW3-94 (red) or JW3-100 (blue), 10 μM raltegravir or 5 μM nevirapine. Mean ± SD (n=2 in technical duplicate). **B** Corresponding induction of IFIT1-luciferase (fold increase above DMSO values) from (A) measured 24 hpi. Unpaired t test, mean ± SD (n=2 in technical duplicate). **C** Corresponding expression from (A) of MxA and IFIT2 (fold induction above DMSO normalised to GAPDH, qRT-PCR 24hpi). Unpaired t test, mean ± SD (n=2 in technical duplicate). **D** Corresponding HIV-1 reverse transcription products from (A) (qPCR 16 hours post infection) shown as % of DMSO- treated control value. Unpaired t-test, mean ± SD (n=2 in technical duplicate). **E** Induction of IRF-luciferase 48 hpi in NT and cGAS -/- THP-1 dual reporter cells following HIV-1 GFP infection and treatment with 1 µM allosteres or DMSO, fold induction compared to uninfected control. Unpaired t test, mean ± SD (n=3 in technical duplicate). **F** Corresponding % infected GFP+ cells from (E) 48 hpi, mean ± SD (n=3 in technical duplicate).

JW3-93 inhibited infection more than JW3-94, despite activating innate immunity less, suggesting that inhibition of RT is the main inhibitory mechanism in these assays, as opposed to innate immune activation (Fig 4A-D). In contrast, in a spreading infection assay with wild type HIV-1 in primary activated T-cells (Fig 1D and E), JW3-94 inhibited infection more effectively than JW3-93, supporting the notion that pharmacological activation of innate immunity can contribute to antiviral effect. Lead allostere JW3-100 also caused innate immune activation in primary monocyte derived macrophages (MDMs) (Supplementary Fig 9D and E), which typically generate a more rapid and potent IFN response than THP-1 cells (Rasaiyaah *et al*, 2013; Sumner *et al*, 2020; Zuliani-Alvarez *et al*, 2022). Importantly, allosteres caused no innate immune activation in the absence of HIV in THP-1 cells (Supplementary Fig 9F).

To examine cGAS involvement in the innate immune activation we observed, we titrated HIV-1 LAI GFP on THP-1 cells encoding an IRF3-sensitive luciferase reporter (THP-1 Dual) with a stable cGAS KO (Supplementary Fig 9G and H), in the presence of 1 µM allostere. As expected, increasing doses of HIV-1 LAI GFP increased luciferase expression in WT cells compared to uninfected cells and addition of 1 µM allostere increased innate immune activation (Fig 4E). Luciferase induction was ablated by cGAS knockout (Fig 4E), consistent with cGAS sensing HIV-1 LAI GFP DNA when allosteres drive premature uncoating of viral capsids. Allostere-inhibited infection was not rescued by cGAS KO, again consistent with innate activation not contributing to antiviral effect in this model (Fig 4F).

### Cofactor binding pocket mutations modulate allostere sensitivity

We hypothesised that designing inhibitors that mimicked cofactors would reduce the opportunity for resistance because resistance mutations should also impact cofactor interactions and reduce viral fitness. However, *in vitro* and *in vivo* studies demonstrate rapid mutation of, otherwise highly conserved (Supplementary Fig 10A), positions in the FG binding site under LEN selection. Mutations include L56I, N57S, M66I, Q67H, Q67Y, K70R, N74D and T107N, with all except L56I and N57S appearing *in vivo* (Yant *et al*, 2019; Link *et al*, 2020; Margot *et al*, 2022a, 2022b, 2023; Segal-Maurer *et al*, 2022). As allosteres are more flexible than LEN, and make fewer interactions across CA monomers (Supplementary Fig 6E) (Bester *et al*, 2020), we hypothesised they may retain activity against LEN resistance mutants. To test this, we infected U87 cells with HIV-1 LAI GFP bearing CA L56I, N57S, M66I, Q67H, Q67Y, K70R, N74D or T107N (Supplementary Fig 10B-C) including lead allosteres (3.2 nM-10 μM) or LEN (3.2 pM-10 nM), and measured infection (GFP expression) and determined IC_50_ (Fig 5A and B). JW3-93/94/100 were broadly inactive against LEN resistance mutations. All three allosteres demonstrated poor activity against HIV bearing L56I or M66I. Structures of allostere-CA complexes indicate that L56 and M66 form hydrophobic interactions with allostere phenyl groups (Supplementary Fig 6C), and so mutation likely weakens binding. The most potent allosteres JW3-93 and JW3-100, but not JW3-94, were less sensitive to N57S with only a 20- and 15-fold increases in IC_50_, respectively, compared to 109-fold for LEN. This may be because there is additional space around N57 with JW3-93/100 complexes compared to JW3-94 as these lack the C2 methyl on the indole. Whilst N57S has not arisen in LEN-treated PLWH (Segal-Maurer *et al*, 2022; Margot *et al*, 2022a, 2022b, 2023), likely due to its essential role in FG-NUP interactions (Dickson *et al*, 2024), this information may be used to guide inhibitor design with reduced resistance sensitivity. Intriguingly, N74D and T107N showed increased infectivity at low allostere concentrations suggesting inhibitor can rescue defective infection by functionally mimicking cofactor binding. We subsequently screened further chemically diverse allosteres, produced in SAR studies, for inhibition of HIV-1 LAI GFP bearing the resistance mutations that arose in LEN-treated PLWH (Supplementary Fig 10D, Table 1 and Supplementary Tables 1-3). However, all eight compounds tested showed similar sensitivities to LEN resistance mutation, suggesting a more extensive SAR campaign is necessary to target capsid inhibitor resistant viruses.

**Figure 5:**
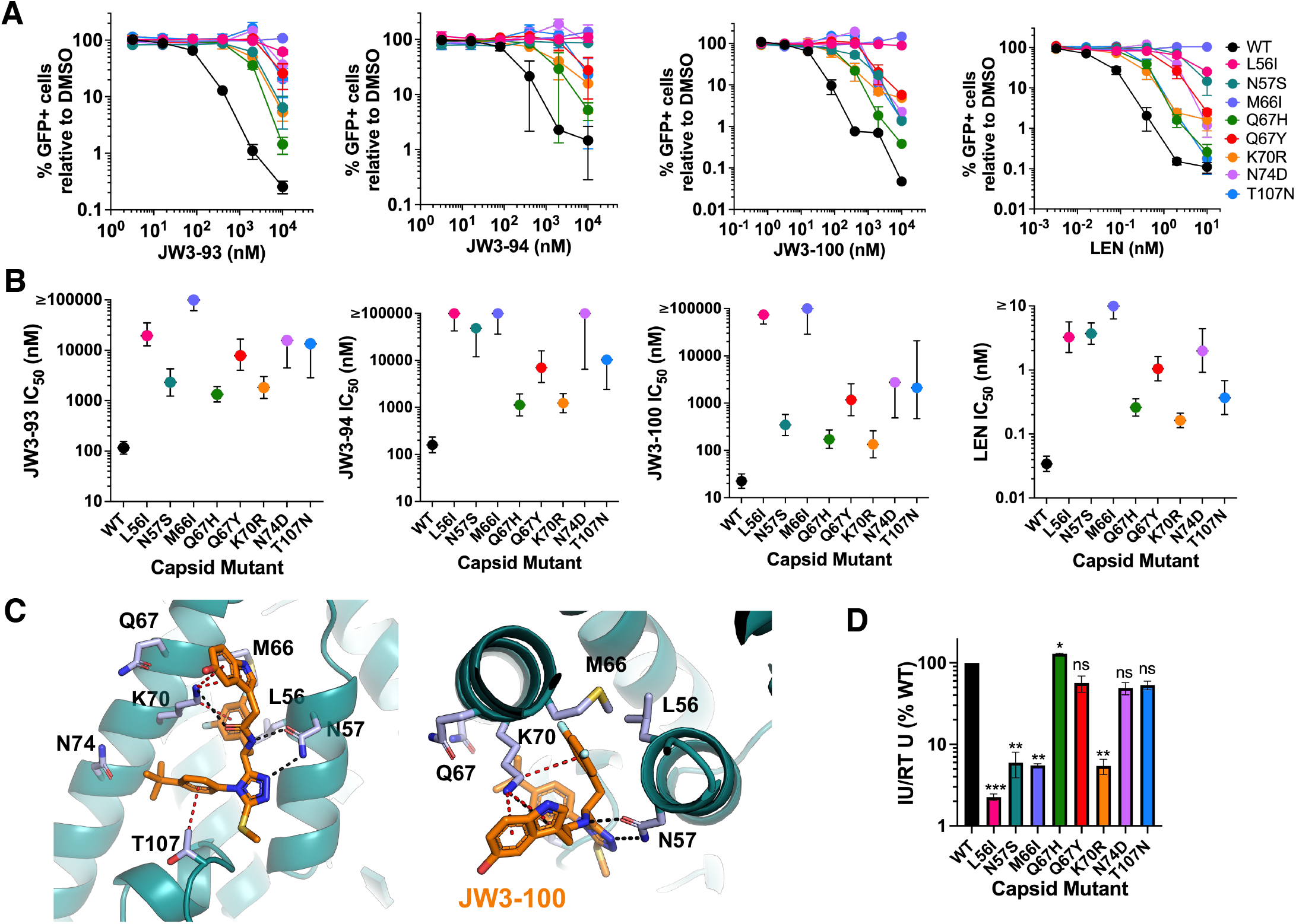
Capsid cofactor binding pocket mutations modulate allostere sensitivity. **A** U87 cell infection with HIV-1 LAI GFP WT and CA mutants (MOI 0.2) with allosteres or lenacapavir treatment. % GFP+ cells (flow cytometry 48 hpi), mean +/- SD (n=2 in technical triplicate). **B** IC50 (95% CI) of JW3-93, JW3-94, JW3- 100 or LEN with HIV-1 GFP mutants calculated by non-linear fit from A. **C** Crystal structure of JW3-100 (orange) bound to CA (PDB-ID 9S6W) with residues where LEN resistance mutations occur coloured in light blue. Black dashed lines: H-bonds/electrostatic interactions. Red dashed lines: π-CH/ π-ion interactions. **D** Infectivity of HIV-1 LAI GFP bearing LEN resistance mutations in U87 cells. Infectious units (IU) calculated using flow cytometry to measure the proportion GFP+ cells following infection with a range of doses of HIV-1 LAI GFP, normalised to reverse transcriptase units (RT U) determined by SG-PERT. P values for each mutant relative to WT by unpaired t-test with Welch’s correction, error bars ± SD (n=2 independent transfections).

Strikingly, predominant LEN resistance mutations Q67H and N74D demonstrated near-WT infectivity in cell lines (Fig 5D) (Yant *et al*, 2019; Link *et al*, 2020) and have appeared rapidly in LEN-treated PLWH (Segal-Maurer *et al*, 2022; Margot *et al*, 2022a, 2022b, 2023; Wirden *et al*, 2024). Q67H arose after only 10 days LEN monotherapy (Margot *et al*, 2022a), whilst, in a separate study, N74D arose after just three weeks of LEN treatment (Wirden *et al*, 2024). Neither Q67 nor N74 directly interact with allosteres (Fig 5C) suggesting a more complex allostere resistance mechanism than inhibitor eviction. Indeed, JW3-93/94/100 all bound cross-linked Q67H hexamers with comparable affinity to WT hexamers (1.5 to 1.8-fold increase in K_D_) by SPR (Table 2) with the small affinity change being unlikely to explain the 7- to 12-fold change in antiviral activity. Similarly, lead allosteres displayed only slight reductions in N74D hexamer binding, with 2.4-3.0-fold increase in K_D_, yet a greater than 100-fold shift in IC_50_. This is in contrast to LEN, where decreased antiviral activity against both Q67H and N74D can be explained by weakened capsid affinity, primarily due to steric hindrance, loss of H-bonds and electrostatic repulsion (Bester *et al*, 2022) (Supplementary Fig 11B-E). These comparisons suggest differences in resistance mechanisms between allosteres and LEN. In the case of allosteres, mutation may confer resistance by interfering with reaction to inhibitor/cofactor binding, either through disrupting the allosteric networks that induce conformational change or through altering capsid stability.

**Table 2:** Affinity (K_D_) of JW3-93, JW3-94 and JW3-100 for WT, Q67H and N74D crosslinked capsid hexamers determined by surface plasmon resonance (SPR). Dissociation constants (K_D_) were calculated by plotting the response at steady state against allostere concentration and performing a sigmoidal fit (n=3 ± SD). P values for each mutant relative to WT by unpaired t-test with Welch’s correction.

|  | WT CA |  | Q67H CA |  |  |  | N74D CA |  |  |  |
| --- | --- | --- | --- | --- | --- | --- | --- | --- | --- | --- |
| Allosteric | $K_D$ (nM) | $IC_{50}$ (nM) | $K_D$ (nM) | Fold change $K_D$ | $IC_{50}$ (nM) | Fold change $IC_{50}$ | $K_D$ (nM) | Fold change $K_D$ | $IC_{50}$ (nM) | Fold change $IC_{50}$ |
| JW3-93 | $165 \pm 60$ | $117 \pm 17$ | $252 \pm 38$ | 1.5<br>($P=0.115$) | $1341 \pm 235$ | 11.5 | $390 \pm 18$ | 2.4<br>($P=0.006$) | >10000 | n/a |
| JW3-94 | $601 \pm 134$ | $159 \pm 30$ | $1057 \pm 298$ | 1.8<br>($P=0.115$) | $1129 \pm 298$ | 7.1 | $1580 \pm 156$ | 2.6<br>($P=0.001$) | >10000 | n/a |
| JW3-100 | $678 \pm 147$ | $23 \pm 4$ | $1129 \pm 101$ | 1.7<br>($P=0.047$) | $172 \pm 36$ | 7.5 | $2014 \pm 40$ | 3.0<br>($P=0.001$) | $2753 \pm 3476$ | 122 |

As previously described (Yant *et al*, 2019; Zhou *et al*, 2015; Shi *et al*, 2015), HIV bearing LEN resistance mutations L56I, N57S, M66I or K70R showed a significant infectivity defect in single round infection in U87 cells, with titres 10- to 40-fold lower than WT virus (Fig 5D). However, despite disruption of key cofactor interactions, and the observed reduction in fitness *in vitro*, mutations M66I and K70R, have arisen in LEN monotherapy treated PLWH at significant frequencies (Segal-Maurer *et al*, 2022; Margot *et al*, 2022a, 2023, 2022b). These observations highlight high CA mutational tolerance and suggest that poor fitness *in vitro* is a poor predictor of whether a virus will be selected and replicate *in vivo*.

### Capsid inhibitor resistance mutants activate innate immunity

Capsid inhibitor resistance mutations are expected to interfere with the cofactor interactions that regulate the timing and location of uncoating. We therefore hypothesized that resistance mutants may inappropriately uncoat, leading to exposure of newly synthesised viral DNA to cGAS and consequent innate immune activation. To test this, we infected THP-1 IFIT1 reporter cells with HIV-1 LAI GFP bearing LEN resistance mutants. As in U87 cells (Fig 5D), there was little difference in the infectivity of Q67H, N74D and T107N-bearing HIV-1 LAI GFP, as compared to WT, in THP-1 IFIT1 cells (Fig 6A). As expected (Sumner *et al*, 2020; Zuliani-Alvarez *et al*, 2022; Cingöz & Goff, 2019), WT HIV-1 LAI GFP induced low-level activation of the IFIT1 luciferase reporter and of MxA, IFIT2 or CXCL10 mRNA expression (qRT-PCR) (Fig 6B and C). In contrast, HIV-1 LAI GFP bearing Q67H, N74D or T107N displayed increased induction of IFIT1, MxA, IFIT2 and CXCL10, as compared to WT, at both 5 and 10 RT U/mL (Fig 6B and C). This suggests that these resistance mutations may compromise capsid integrity via intrinsic destabilisation or through preventing appropriate interactions with, and/or responses to, the cofactors that regulate capsid stability. Crucially, this demonstrates that enhanced innate immune triggering does not prevent replication *in vivo*, as these mutations have arisen in LEN treated PLWH.

**Figure 6:**
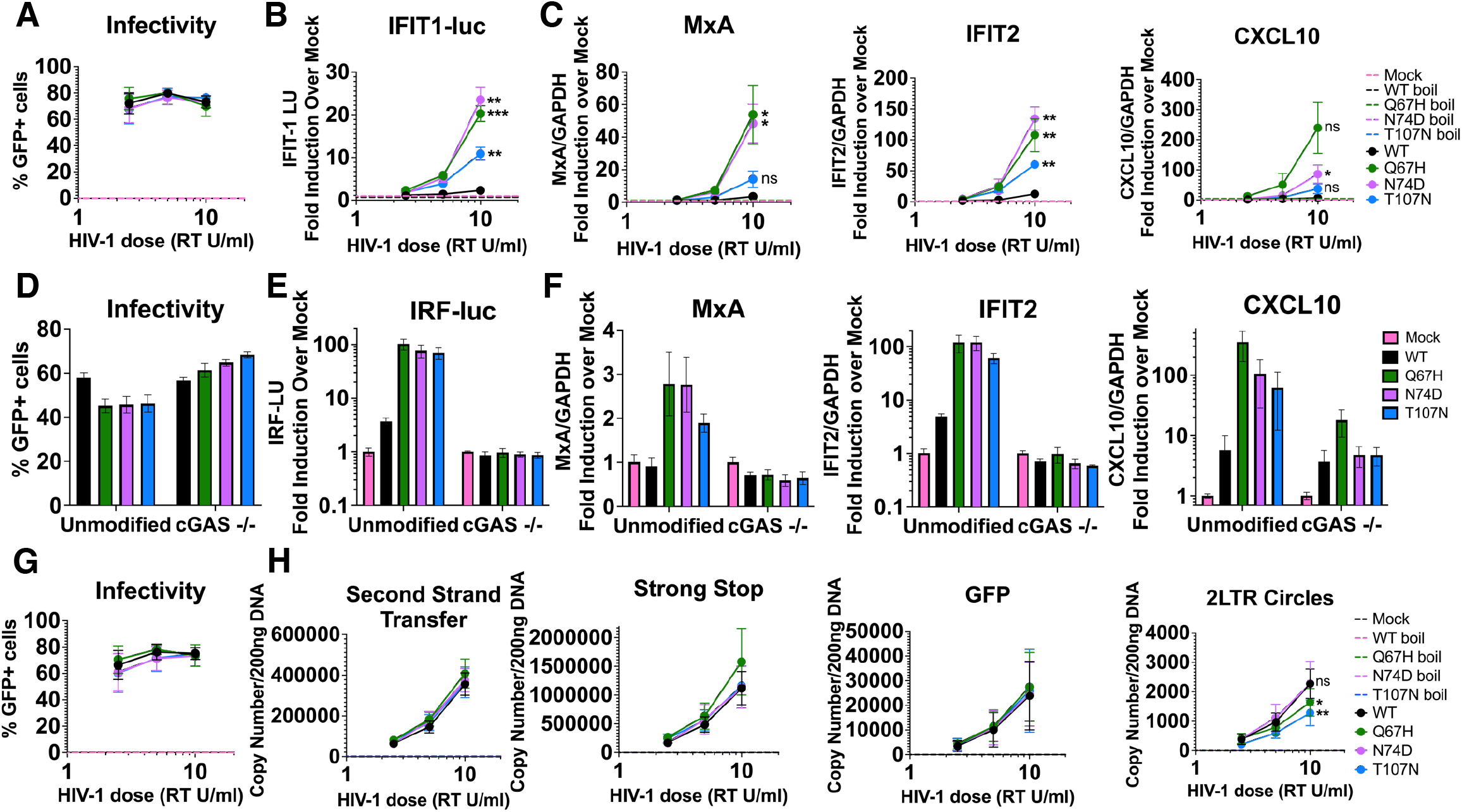
HIV-1 bearing lenacapavir resistance mutations activates innate immunity without inhibitor. **A** % GFP+ THP-1 IFIT1 cells following infection with HIV-1 LAI GFP WT and CA mutants (2.5-10 RT U/ml). (flow cytometry 48 hpi) ± SD (n=2 in technical duplicate). **B** Corresponding induction of IFIT1-luciferase for (A) shown as fold increase compared to uninfected cells (luminometry 48 hpi) ± SD (n=2 in technical duplicate). **C** Corresponding expression for (A) for MxA, IFIT2 and CXCL10 shown as fold induction over GAPDH and uninfected values (qRTPCR 24 hpi) ± SD (n=2 in technical duplicate). **D** Proportion of GFP+ THP-1 IFIT1 cells following infection with HIV-1 LAI GFP WT and CA mutants (2.5-10 RT U/ml) (flow cytometry 48 ± SD (n=3 in technical duplicate). **E** Corresponding levels of HIV-1 reverse transcription products for (D): strong stop, second strand transfer, GFP and 2-LTR circles (qPCR 16 hpi). Unpaired t-test. Error bars ± SD (n=3 independent experiments in technical duplicate). **F** Proportion of GFP+ THP-1 Dual reporter cells following infection with HIV-1 LAI GFP WT and CA mutants (5 RT U/ml). Measured by flow cytometry 48 hours post infection. Error bars ± SD (n=2 independent experiments in technical duplicate). **G** Corresponding induction of IRF-luciferase shown as fold increase compared to uninfected cells. Measured by luminometry 48 hours post infection. Error bars ± SD (n=2 independent experiments in technical duplicate). H Corresponding expression of MxA, IFIT2 and CXCL10 shown as fold induction compared to GAPDH levels and uninfected cells. Measured by qPCR 24 hpi. Error bars ± SD (n=2 in technical duplicate)

Notably, the HIV mutants did not induce IRF luciferase reporter or ISG expression in cGAS KO cells (Fig 6D-F). This was not a result of reduced infection by the mutants in the cGAS KO cells, because they were slightly more infectious than WT virus, suggesting a small cGAS dependent inhibition of mutant HIV infection in these cells (Fig 6D). If the LEN resistance mutations caused premature uncoating, they may also affect RT (Papa *et al*, 2023; Rebensburg *et al*, 2021). However, we found that HIV-1 LAI GFP with Q67H, N74D and T107N generated equivalent levels of early (strong stop) and late (second strand transfer and GFP primers) products to WT virus (Fig 6H), consistent with these mutations having minimal impact on infection (Fig 6G) and, ruling out increased amounts of DNA as the explanations for increased innate immune activation. This suggests that, unlike allosteres, these mutations do not completely compromise capsid core integrity, which is required for efficient RT (Christensen *et al*, 2020; Jennings *et al*, 2020; Sowd *et al*, 2021), but cause a more subtle defect that exposes the genome but retains normal levels of DNA synthesis. A small reduction in 2-LTR formation (nuclear import) was observed for the Q67H and T107N mutants (Fig 6E). This is expected given Q67 and T107 contact NUP153 and CPSF6, both of which are cofactors for nuclear transport. There was no reduction in 2-LTR circles with N74D infection, thought to be because this mutant interacts effectively with NPC FGs, but fails to interact with CPSF6 and disengage from the pore, and simply integrates into chromatin that comes close enough to the NPC (Bejarano *et al*, 2019; Zila *et al*, 2019).

## Discussion

Here we describe an HIV capsid-targeting inhibitor series with nanomolar potency, developed by designing molecules that mimic the FG-bearing host cofactors that regulate HIV capsid stability and uncoating location. Solving structures of native CA hexamer-allostere complexes, and comparing with hexamer-FG cofactor peptide complexes, revealed that inhibitors cause allosteric shifts in the hexamer CA-CTD at the lattice two- and three-fold symmetry axes, thereby altering the mechanical behaviour of the capsid during specific infection stages. Notably, host cofactors elicit distinct, localised structural responses that facilitate the mechanical requirements of the viral lifecycle. NUP153 markedly stabilizes the two-fold interface -featuring a highly specific interaction (−9.8 kcal/mol), which likely provides the local stability required for nuclear pore translocation. Conversely CPSF6 acts as a “molecular glue” at the three-fold vertices, doubling the interface area compared to the apo form to lock the global fullerene lattice together.

In contrast, our allosteric inhibitors occupy the shared FG-binding pocket but fail to induce these critical cofactor driven effects. Specifically, they do not provide the robust three-fold reinforcement seen with CPSF6, nor the specific two-fold stabilization of NUP153. We hypothesize that natural HIV uncoating is driven by a combination of the additive allosteric effects of cofactor recruitment, where the hand-off from NUP153 to CPSF6 transitions the capsid from a flexible state (optimised for pore transit) to a rigid, “locked” state (optimised for intra-nuclear transport to chromatin). This transition likely occurs with the internal pressure changes associated with DNA synthesis completion (Burdick *et al*, 2024; Müller *et al*, 2021).

In our proposed model, allosteres prevent NUP153 and CPSF6 binding and, while they stabilize individual hexamers, they limit the CA lattice’s mechanical adaptability. By slightly rigidifying the hexamer-hexamer edges (two-fold) while failing to reinforce the three-fold vertices, these inhibitors promote a “brittle” lattice. Such a structure may fail to withstand the mechanical stresses normally experienced by the virus and particularly during nuclear transport. Furthermore, this rigidified hexameric geometry may be incompatible with the highly curved regions of the capsid where pentamers are concentrated. If inhibitors favour hexameric packing at the expense of pentamer integration, the resulting lattice would be prone to rupture at “cap-like structures” (Rankovic *et al*, 2018). This mechanism may explain why particles produced in the presence of capsid inhibitors PF74 and LEN exhibit malformed, hyperstable, yet non-infectious capsid assemblies (Link *et al*, 2020; Faysal *et al*, 2024).

CA residue K182 is a key residue in the allosteric cascade, located at the N-terminal of H9 α-helix. We show that all examined inhibitors, including PF74 and LEN, promote a shared conformational profile: subtle shifts in helices H8 and H9 at the two-fold edges and a consistent shift in the C-terminal helices H10 and H11 that locks the three-fold corner into a structurally weak interaction. The most striking feature of our series is the specific H-bonding of inhibitors JW3-93 and JW3-100 with K182. Because K182 serves as a non-mutable linchpin for the H9-H11 relay, the virus cannot easily mutate this residue to escape the inhibitor without compromising the fundamental stability of the capsid lattice (Saito *et al*, 2019). This makes the K182-hydroxyl-indole interaction a potent site for resisting viral resistance mutations, effectively “trapping” the capsid in a state that is functionally incompatible with the host-factor-mediated uncoating program.

HIV capsid inhibitor dose-response curves are typically bi- or triphasic, likely representing different inhibition modes. The first phase may represent low occupancy inhibition of FG-cofactor interactions, suppressing timely regulation of uncoating and nuclear transport. A second phase, requiring higher occupancy, is thought to represent catastrophic uncoating with consequent inhibition of DNA synthesis (Márquez *et al*, 2018; Price *et al*, 2014; Saito *et al*, 2016). We found that JW3-94 exhibits a triphasic dose-response curve consistent with different modes of action at low and high concentrations. JW3-93 however, has a monophasic dose-response curve, suggesting a single mode of action (Fig 1), likely related to capsid uncoating, evidenced by RT inhibition at low dose. This is likely due to the indole group, the only structural difference from JW3-94, which makes additional H-bonds across the monomer-monomer interface (Fig 3). Molecules forming more cross-monomer interactions should have a greater stabilising, and thus flattening, effect on capsid lattices and are expected to break capsids open at lower occupancy (Faysal *et al*, 2024). Thus, JW3-93 may rupture capsids at lower occupancy than JW3-94, consistent with JW3-93 causing inhibition of RT at lower doses than JW3-94 (Fig 4). Uncoating cores at lower concentration may mask effects of inhibition of cofactor binding leading JW3-93 inhibition to be predominantly mediated by breaking capsids open prior to DNA synthesis.

Strikingly, CypA loss shifted the upper part of the JW3-100 inhibition curve to the right. This effect has previously been observed for inhibition of HIV by PF74 (Zhou *et al*, 2015; Saito *et al*, 2016; Shi *et al*, 2011) and suggests that in the absence of CypA, inhibition of cofactor binding by low-dose inhibitor no longer contributes to inhibitory effect. This is consistent with a role for CypA impacting cofactor activity by allosterically regulating conformation and/or dynamics at the FG-binding site (Twarock *et al*, 2024; Morling *et al*, 2025). Notably, this allosteric model is also supported by magic angle spinning NMR experiments which showed chemical shift perturbations at FG cofactor-binding CA N57 upon CypA recruitment (Lu *et al*, 2015). Functionally, this allosteric model is also supported by the observation that CypA inhibition renders HIV insensitive to downstream FG cofactor depletion (Schaller *et al*, 2011).

Our data support a model in which allosteres activate innate immune sensing against infection by breaking capsids open prematurely, i.e. at the wrong time, in the wrong place, revealing newly formed viral DNA to cGAS. Similar *in vitro* observations have been made for PF74 (Sumner *et al*, 2020; Kumar *et al*, 2018) and low dose LEN (Eschbach *et al*, 2024; Scott *et al*, 2025), as well as with other methods of disrupting capsid integrity including HIV protease inhibitors, Gag cleavage mutants and mutants that prevent IP6 packaging (Sumner *et al*, 2020; Papa *et al*, 2023). This suggests that capsid inhibitor potency might be enhanced *in vivo* by natural innate immunity and interferon production activated downstream of cGAS sensing.

We find that mutations promoting allostere and LEN resistance (Fig 5) cause HIV to activate cGAS and subsequent interferon-sensitive gene expression *in vitro* in the absence of inhibitor (Fig 6). We expected innate immune activating mutations would not be selected *in vivo* because immune activation should limit replication, particularly *in vivo*. However, LEN monotherapy rapidly selects CA mutants (Segal-Maurer *et al*, 2022; Margot *et al*, 2022a, 2022b, 2023), that trigger innate sensing *in vitro* eg N74D (Rasaiyaah *et al*, 2013), and Q67H and T107N (Fig 6) suggesting innate immune activation is not a barrier to replication once infection is established. However, we hypothesise that innate immune activation reduces transmission frequency. This is evidenced by our observation that non-pandemic HIV-2 and HIV-1(O), which are relatively poor transmitters, are better activators of innate immune sensing by cGAS and TRIM5, than the best transmitter - pandemic HIV-1(M). Pandemic HIV adaptations include a more dynamic capsid surface suggesting that increased CA dynamics facilitate a more precise regulation of Capsids to prevent premature uncoating and DNA release. SIV transmission studies in macaques also support an important role for innate immunity in transmission - IFN effectively inhibited transmission and interferon inhibition enhanced transmission (Sandler *et al*, 2014). Thus, capsid inhibitors, that expose viral nucleic acid might reasonably be expected to be most useful as pre-exposure prophylaxis (PrEP), where innate immune activation is expected to contribute to prevention of transmission. Indeed, LEN is currently undergoing clinical trial (PURPOSE) for use as an injectable monotherapy PrEP and has been approved as PrEP by the FDA, where its long-acting nature is expected to mitigate the inconvenience and stigma of daily pills (Bekker *et al*, 2024).

LEN resistance mutations triggering innate sensing may also contribute to LEN effectiveness in PrEP. Typically, combination therapy is necessary to avoid development of drug resistance, even for prophylaxis (Tang & Shafer, 2012). For example, the very effective integrase strand transfer inhibitors (INSTIs) including raltegravir, dolutegravir and long-acting cabotegravir (CAB-LA) have all been trialled as monotherapies as ART and PrEP, with a view to simplifying inhibitor regime (Loosli *et al*, 2023; Koss *et al*, 2024; Blanco *et al*, 2018; Caby *et al*, 2010). However, resistance mutations arose with resistance also detected in treatment-naive PLWH, suggesting transmission (Hao *et al*, 2025; Šablinskaja *et al*, 2025; Mielczak *et al*, 2024; Geremia *et al*, 2024). Importantly, CAB-LA as PreP has led to breakthrough transmission and subsequent selection of resistant virus. Our demonstration of LEN resistant mutants activating cGAS, together with evidence for effective innate immune inhibition of transmission, suggests that LEN resistant mutant transmission should be rare. This notion is supported by 2 clinical trials. In 2134 cisgender women treated with LEN at 26-week intervals there were no breakthrough infections (Bekker *et al*, 2024). In PURPOSE 2179 men and gender-diverse persons were similarly treated with LEN, and in this case, 2 participants in the LEN group had acquired HIV at the time of primary analysis (Kelley *et al*, 2025). Virus from both positive participants bore CA N74D at diagnosis. It is not clear whether virus bearing N74D was transmitted or whether effective LEN monotherapy selected for CA N74D after transmission of wild type virus, but the latter explanation seems most likely.

CA inhibitor derivatives with increased resistance to escape have been considered. Increasing the flexibility of the LEN cyclopenta-pyrazole ring, which sits close to H67, through replacement with a tetrahydroindazole ring, removing the cyclopropyl moiety and two fluorine atoms, increases potency against WT HIV-1 and reduces sensitivity to the Q67H mutation (Bester *et al*, 2022). Future work might focus on modifying allosteres to retain activity against such mutations by increasing inhibitor flexibility, for example by adding alkane chains or remodelling the triazole group, to enable the molecule to retain binding to a mutated CA pocket. Forming additional interactions with the CA pocket in regions distant from the residues prone to mutation would also increase affinity and could prevent inhibitor eviction by pocket mutation. In addition, targeting alternative cofactor binding sites on the capsid, such as the central hexameric pore, or the three-fold symmetry axis in the capsid lattice, remain underexplored approaches likely to be effective. Certainly, the commitment to drugging HIV capsid has demonstrated the tractability of targeting viral capsids and mimicking host-cofactor interactions. Of course, all viruses have host co-factors and we argue that the development of LEN, and the work herein, demonstrates clearly how a sound understanding of virus-cofactor interactions and their function can drive therapeutic innovation. Indeed, we propose the effectiveness of LEN in HIV prophylaxis, with the contribution of innate immune sensing to its unique mechanism, demonstrates how such new knowledge can transform antiviral development, and in this case, provide a realistic and tractable tool for HIV eradication.

## Methods

### Methods and Protocols

#### Chemical synthesis and compound characterisation

Synthetic methods and characterisation for lead compounds are provided in the supplementary information. All compounds were dissolved in anhydrous DMSO (900645, Sigma Aldrich) and stored in screw cap tubes with drying agent.

Compound stability assays in hepatocytes were performed as described in (Smith *et al*, 2022).

#### Protein production and purification

Cross-linked and non-cross-linked HIV-1 CA was expressed and purified as previously described (Gres *et al*, 2015; Price *et al*, 2014). In brief, the plasmids encoding cross-linked or non-cross-linked CA were transformed into *E. coli* OverExpress™ C41(DE3) (Lucigen). Single colonies were used to inoculate overnight starter cultures of LB medium (100 μg/mL ampicillin). The following day this was used to inoculate 1 L LB medium (100 μg/mL ampicillin) and grown at 37°C in an orbital shaker at 250 rpm until the OD600 nm reached 0.5. Overexpression was induced by adding IPTG to a final concentration of 0.4 mM and protein expression was carried out overnight at 14°C. Cells were harvested by centrifugation at 6000 g for 20 min. The pellets were resuspended in 50 mL lysis buffer (50 mM Tris-HCl, 40 mM NaCl, 20 mM β-mercaptoethanol (βME) [pH 4.5]) per 1 L culture and lysed by sonication. The lysate was cleared by centrifugation at 30000 g for 20 min. To precipitate the capsid protein the cleared lysate was subsequently added ammonium sulphate to a final concentration of 20 % (w/v) and pelleted by centrifugation at 30000 g for 20 min. Pellets were resuspended in 10 mL refolding buffer (100 mM citric acid, 20 mM βME [pH 4.5]) per L culture and after an extensive dialysis against the same buffer any remaining insoluble material was removed by centrifugation.

After the initial refolding non-cross-linked capsid were dialysed directly against 25 mM Tris-HCl [pH 8] before being further purified by Anion Exchange Chromatography (AEC) using 5 mL Hi-TRAP Q columns (Cytiva). AEC was performed using Buffer A (25 mM Tris-HCl [pH 8]) and Buffer B (25 mM Tris-HCl, 1M NaCl [pH 8]). Lastly Size Exclusion Chromatography (SEC) was performed using a Superdex 16/600 75 pg column (Cytiva) with a buffer containing 25 mM Tris-HCl and 40 mM NaCl [pH 8]. All steps were performed on an ÄKTA pure system (Cytiva). The protein was further concentrated to 3.0 mg/mL in the SEC buffer for crystallisation. Cross-linked capsid protein were dialysed in three steps (Step 1 (50 mM Tris-HCl, 1 M NaCl, 20 mM βME, [pH 8]), Step 2 (50 mM Tris-HCl, 1 M NaCl, [pH 8]), (50 mM Tris-HCl, 40 mM NaCl, [pH 8]) to facilitate the cysteine cross-linking. AEC was performed similarly as with non-cross-linked capsid, but with 40 mM NaCl added to Buffer A. SEC was performed using a Superdex 16/600 200 pg column (Cytiva).

#### Crystallisation, structure solution and analysis

Crystals were grown using the hanging-drop vapour diffusion technique at 20°C by mixing 1 μL protein with 1 or 2 μL of precipitant, containing 9.5-11% (w/v) PEG3350, 250-350 mM NaI, and 100 mM Sodium Cacodylate, [pH 6.5] as previously described (Gres *et al*, 2015). Crystals of hexagonal shape appeared after 3-5 days reaching a maximum size of 0.1 – 0.2 mm after 2 weeks. At this stage they were soaked with 0.5-1 mM of each allostere or peptide dissolved in DMSO, and incubated at 20°C for 45-180 mins before crystal harvesting. The harvested crystals were immersed in a solution containing the precipitant mixture and 20 % (v/v) glycerol and cryo-cooled in liquid nitrogen. Diffraction data were collected from single crystals at the PETRA III P13 or the P14 beamline (EMBL-Hamburg/DESY P13, Germany), and ID23-1 (ESRF, Grenoble, France) and I04 (Diamond Light Source, Oxfordshire, UK). All data sets were indexed, processed, and scaled using XDS (Kabsch, 2010) or autoPROC (Vonrhein *et al*, 2011). All HIV-1 (M) CA crystals belonged to the P6 space group with a solvent content of 48.5% corresponding to one molecule per asymmetric unit and consistent cell volumes across crystal structures which vary only ∼0.5% with no correlation to any specific bound ligand (Supplementary Table 6).

The structures were determined by molecular replacement using Phenix Phaser (Adams *et al*, 2010) with a previously determined HIV-1(M) CA structure (PDB ID:4XFX) as a search model. Model building was performed using COOT (Emsley & Cowtan, 2004). Refinement was performed using BUSTER (Vonrhein *et al*, 2024), REFMAC 5.8 (Murshudov *et al*, 2011) or Phenix refine (Afonine *et al*, 2012) using a TLS/maximum likelihood protocol.

The CPSF6 peptide (Pro313-Gly327) was modelled in the CA complex from Pro313 to Pro324 while the NUP153 peptide (residues Thr1407-Thr1423) was modelled from Asn1409 up to Gly1418. CA in the CA-NUP153 complex was modelled up to Gln219 (main chain atoms only) because beyond this it was disordered. For all the other structures there was clear electron density up to residue Val221. Both peptide complexes closely resemble the previously published structures determined in native background (PDB IDs 6AY9 and 6AYA) (Gres *et al*, 2023) with RMSDs 0.484 Å and 0.379 Å respectively.

Mature CA hexamers could be assembled by crystallographic symmetry operators. Based on difference anomalous Fourier maps, we have confirmed the presence of two iodine ions in the asymmetric unit, one coordinated by Arg173 and Lys170 and a second on the loop Gln155-Glu159 with both conserved in all previous uncross-linked structures. Additional peaks in the electron density maps were modelled as Chlorine or Sodium atoms based on the chemical environment and geometry around them. We assume that the iodines, and to a lesser degree the other atoms, contribute to stabilising the hexamers in the absence of cross-links.

Final figures were rendered in The PyMOL Molecular Graphics System (Schrödinger, LLC.™). Structural alignments and protein interfaces were performed using the EBI servers PDBeFold (Krissinel & Henrick, 2004) and PDBePISA (Krissinel & Henrick, 2007).

#### Surface Plasmon Resonance (SPR)

SPR experiments were performed at 25 °C using a Biacore T200 with CM5 sensor chips (Cytiva). Cross-linked CA hexamers were immobilised in PBS-P+ buffer by flowing 1:1 N-hydroxysuccinimide (NHS) and N-(3-dimethyl-aminopropyl)-N’-ethyl-carbodiimide hydrochloride (EDC) for 420s at 10 µL/min, followed by CA at 100 μg/mL in 10 mM sodium acetate [pH 5] for 420s, then ethanolamine (1M [pH 8]) for 420s. Kinetics/affinity experiments were performed in 10 mM HEPES [pH 7.4], 150 mM NaCl, 3 mM EDTA, 0.05% P20 (HBS-EP+) with 5% DMSO. Inhibitors (0.195 µM-25 µM) were generally injected for 120s followed by 300s dissociation. No regeneration step was used. Dual flow cells were used with a blank reference with no protein injection. Solvent correction was carried out using 4-6% DMSO. Each allostere concentration was run in technical duplicate and a wash of 50% DMSO 50% HBS-EP+ was performed following each injection.

#### Cell lines

CCR5 expressing U87 (ATCC) and HEK 293T (ATCC CRL-3216) cells were maintained in Dulbecco’s Modified Eagle Medium (DMEM) (Gibco, 41966-029), with 10% fetal calf serum (FCS, LabTech), 100 U/mL penicillin and 100 μg/mL streptomycin (Pen/Strep, Gibco, 15140-122) at 37 °C in 5% and 10% CO_2_, respectively. THP-1 cells were maintained at 2×10^5^ cells/mL in Roswell Park Memorial Institute medium (RPMI) (Gibco, 21875-034), 10% FCS and Pen/Strep at 37 °C in 5% CO_2_. THP-1-IFIT1 cells which express Gaussia luciferase under control of the IFIT-1 promoter were previously described (Mankan *et al*, 2014). THP-1 Dual cells (cGAS KO and NT control, Invivogen) express Lucia luciferase as a marker of the IRF pathway and were additionally supplemented with 12.5 mM HEPES (Sigma), 10 μg/mL of blasticidin (Invivogen) and 100 μg/mL of zeocin (Invivogen).

#### Isolation of primary CD4+ T cells from peripheral blood

Peripheral blood mononuclear cells (PBMC) were isolated from leukocyte cones from healthy donors (UK NHS Blood and Transplant Service) by density centrifugation using FicollPaque Plus (GE Life Sciences). Resting CD4+ T cells were isolated from total PBMCs by negative selection using the CD4+ T cell Isolation Kit (130-096-533, Miltenyi Biotec). T-cells were cultured in RPMI 1640 with 10 % FBS and 10 IU/mL IL-2 (Centre For AIDS Reagents (CFAR), National Institute of Biological Standards and Control (NIBSC), cat# 0901). Following isolation, T cells were stained with CD3-APC (981012, Biolegend) and CD4-PE (980804, Biolegend) and analysed by flow cytometry to check purity (≥ 97 % CD3+ CD4+ for all donors). T cells were activated for 5 days in culture medium in the presence of 1 µg/mL plate-bound anti-CD3 antibody (clone OKT3, 16-0037-85, Thermo Fisher Scientific) and 2 µg/mL soluble anti-CD28 antibody (clone CD28.2, 16-0289-85, Thermo Fisher Scientific).

#### Isolation of primary monocyte-derived macrophages

Monocyte-derived macrophages (MDMs) were purified from PBMCs isolated from fresh blood from healthy volunteers, as approved by the joint University College London/University College London Hospitals NHS Trust Human Research Ethics Committee, and written informed consent was obtained from all participants. PBMCs were washed three times in PBS and adherent cells selected by plating for 1.5 hours before washing. These were then incubated in RPMI with 10% heat-inactivated pooled human serum (Sigma) and 40 ng/mL macrophage colony-stimulating factor (R&D systems) for 3 days before replacing the media with RPMI with 10% FBS and incubating for a further 3-4 days prior to infection.

#### Generation of LEN resistance mutations in pLAIΔEnv GFP, pOPT HIV-1 (M) Hex and pOPT HIV-1 (M)

pLAIΔEnv GFP, pOPT HIV-1 and pOPT HIV-1 Hex containing LEN resistance mutations were generated by site-directed mutagenesis using Pfu Turbo DNA polymerase (Agilent, 600250) and the primers below. For pLAIΔEnv GFP, part of the capsid gene was first TOPO cloned using the primers described in reagents table, then excised by restriction digest with BSSHII (NEB, R0199S) and AgeI-HF (NEB, R3552S) and cloned back into pLAIΔEnv GFP.

#### Production of single round infection HIV-1 GFP and HIV-1 LAI GFP

Sub-confluent HEK 293T cells in 10 cm dishes were transfected with 1.5 μg pCSGW, 1 μg p8.91 and 1 μg pMDG (HIV-1 GFP), or 2.5 μg pLAI and 1 μg pMDG (HIV-1 pLAI ΔEnv GFP) with 200 μL Opti-MEM (Gibco, 31985070) and 10 μL FuGENE 6 transfection reagent (Promega, E2692), according to the manufacturer’s protocol. Alternatively, 8×10^5^ cells were seeded onto 6-well plates and a third of the plasmids and transfection reagents used. Media was changed after 24 hours. Supernatants were harvested 48- and 72-hours post-transfection and purified through 0.45 μm syringe filters. For innate immune sensing assays and when measuring HIV-1 reverse transcripts, virus preps were treated with 20 U/mL deoxyribonuclease (DNase) 1 (Sigma, DN25) with 10 mM MgCl_2_ for 2 hours at 37°C. Where necessary to achieve desired multiplicity of infection (MOI), virus was concentrated by ultracentrifugation at 23000 rpm for 2 hours and 4°C through 20% sucrose (Thermo Scientific Sorvall WX+ Ultra Series) and resuspended in RPMI media. Preps were aliquoted and stored at -80°C.

Reverse transcriptase activity was quantified using SYBR Green PCR enhanced RT assay (SG-PERT) as previously described (Vermeire *et al*, 2012; Pizzato *et al*, 2009). A QuantStudio5 (Thermo Fisher Scientific) was used for qPCR. Results were analysed using Thermo Fisher Design and Analysis software. To determine multiplicity of infection (MOI), U87 cells were seeded 24 hours prior to infection at 5×10^4^ cells/well in a final volume of 250 μL in a 48-well plate or 1×10^5^ cells/well in a final volume of 2 mL in a 6-well plate. THP-1 cells were seeded at the time of infection at 1.25×10^4^ cells/well in a final volume of 250 μL in a 48-well plate. Cells were infected with a range of doses of virus and 8 μg/mL polybrene (Sigma), incubated for 48 hours, fixed with 4% paraformaldehyde and the proportion of GFP positive cells measured by flow cytometry as described.

#### Production of full-length HIV-1 NL4-3

The HIV-1 clone pNL4-3 was obtained from the CFAR, NIBSC (2006). NL4-3 stocks were produced by plasmid transfection of HEK 293T cells with Fugene 6 (Promega, E2692). Supernatants were harvested at 48 hrs and 72 hrs, filtered, DNase treated, purified and concentrated by ultracentrifugation through a 25% sucrose cushion and resuspended in RPMI 1640 with 10% FBS. Viral titres were determined by measuring reverse transcriptase activity in mU by SG-PERT assay.

#### Allostere virus production assays

HEK 293T cells were seeded in 6-well plates at 8×10^5^ cells/well. 24 hours later, they were transfected with 750 ng pLAI and 300 ng pMDG in 60 μL Opti-MEM and 3 μL FuGENE 6 (Promega, E2692). 18 hours post-transfection, media was changed and 5 μM allosteres added. 48-hours post-transfection cells and supernatants were collected. RT activity was quantified by SG-PERT. p24 levels in the cell extracts and supernatants was quantified by western blot with actin and VSV-G loading controls, respectively. To measure infectivity, supernatants were diluted 1000-fold and a range of doses used to infect U87 cells seeded at 1.25×10^4^ cells/well in 48-well plates 24 hours prior.

#### HIV-1 infection to measure allostere inhibition

For single-round infection, U87 cells were seeded at 1.25×10^4^ cells/well in 250 μL in 48-well plates or 1×10^5^ cells/well in a final volume of 2 mL in 6-well plates. After 24 hours, cells were infected with HIV-1 GFP or HIV-1 LAI GFP (MOI 0.3) and treated with 0.313-10 μM compounds and 8 μg/mL polybrene.

To infect activated primary CD4+ T cells with HIV-1, cells were mixed and incubated with 200 mU reverse transcriptase of HIV-1 NL4-3 per 10^6^ cells for 4 hrs at 37°C. T cells were subsequently washed twice in PBS by centrifugation (400g, 5 mins) and resuspended in fresh culture medium. At the indicated time points, cells or culture supernatants were harvested to determine intracellular infection levels by Gag staining or virus release by SG-PERT, respectively. Treatment with the indicated compounds (500 nM) or DMSO was maintained throughout the experiment. The experiment was carried out in duplicate for each donor. Cells were supplemented with fresh IL-2 four days post infection.

#### Flow cytometry

For U87 and THP-1 cell line experiments, 48 hours post-infection, cells were trypsinised, fixed with 4% paraformaldehyde in PBS and the proportion of GFP positive cells measured by flow cytometry using a BD Accuri C6 machine, FACS calibur machine (BD Bioscience) or NovoCyte (Agilent). Experiments were performed in technical triplicate with uninfected negative controls. Data were analysed on NovoExpress 1.5.0 software (Agilent).

For primary T cell infection experiments, cells were washed in PBS and stained with fixable Zombie R685 Live/Dead dye (423119, Biolegend, 1:500) for 5 mins at 37°C. Excess stain was quenched with FBS-complemented RPMI. Cells were fixed with 4% formaldehyde before intracellular staining. Permeabilisation for intracellular staining of primary T cells was performed with Permeabilisation Wash Buffer (421002, Biolegend) according to the manufacturer’s instructions. Intracellular staining for HIV-1 Gag (KC57-FITC, clone FH190-1-1, 6604665, Beckman Coulter, 1:100) was performed for 30 mins at room temperature followed by washing with PBS (1700 rpm, 5 mins, 4°C) to remove excess antibody. Data were acquired on a NovoSampler Pro (Agilent) and analysed using NovoExpress 1.5.0 software (Agilent).

#### MTT assay for cell viability

U87 cells were seeded at 5×10^3^ cells/well in 100 μL in 96-well plates. After 24 hours, cells were treated with 10 μM allosteres. After 48 hours, 10% v/v 3-4-5-dimethylthiazol-2-yl-2,5-diphenyltetrazolium bromide (MTT, Sigma) was added and incubated for 2 hours. Resulting crystals were solubilised with 100 μL 10% SDS with 0.01M HCl for one hour. Absorbance was measured at 570 nm using a platereader (Thermo Scientific MultiskanTM FC Microplate Reader) and results were normalised to background wells with no cells.

#### Luciferase reporter assays for innate immune activation

THP-1 IFIT1 or dual cells at 7.5×10^5^ cells/well, DNase-treated HIV-1 LAI GFP (MOI 0.4), 0.1-10 μM allosteres and 8 μg/mL polybrene were combined in 6-well plates in a final volume of 1.5 ml. After 24 and 48 hours, 10 μL of supernatant was added to a white 96-well plate. Gaussia luciferase renilla substrate coelenterazine (NanoLight Technology, 303-500) was diluted to 1 mg/mL in ethanol then further diluted 1:500 in PBS. 50 μL was added to the supernatant using automated injectors and luciferase read out using a GloMax Navigator luminometer (Promega). Fold induction of IRF3/IFIT1 was normalised to uninfected or DMSO treated cells.

#### Quantitative PCR (qPCR) to measure RT products and ISGs

THP-1 IFIT1 cells at 7.5×10^5^ cells/well in 6-well plates, were treated with 0.1-10 μM allosteres and infected with DNase-treated HIV-1 LAI GFP (MOI 0.4). To measure the effect of CA mutants, THP-1 IFIT1 cells were seeded at 7.5×10^5^ cells/well in 6 well plates in a final volume of 1.5 ml, treated with 8 μg/mL polybrene and infected with 2.5, 5 and 10 RT U/mL DNase-treated and sucrose-purified HIV-1 LAI GFP with CA WT, Q67H, N74D and T107N. Alternatively, cells were seeded at 3×10^5^ cells/well in 24 well plates in a final volume of 600 μL. Virus which had been boiled for 10 minutes was included as a control for plasmid in the virus prep. Each condition was set up in duplicate. 16 hours post infection samples were taken to measure RT products, 24 hours post infection samples were taken to measure ISGs and IFIT1 luciferase as above and at 48 hours post infection samples were taken to measure IFIT1 luciferase and the proportion of GFP positive cells by FACS as above.

To measure RT products, DNA was extracted from cells using DNeasy Blood and Tissue Kit (Qiagen 69506), according to the manufacturer’s protocol. DNA concentration was normalised using NanoDrop. TaqMan qPCR was used to quantify strong stop, second strand transfer, GFP and 2LTR circles using primers detailed below. Reactions were set up in 384-well plates with 5 μL TaqMan Gene Expression Master Mix (Applied Biosystems 4369016), 0.2 μL 10 μM FAM/TAMRA probe, 1 μL 10 μM each forward and reverse primers, 0.8 μL dH_2_O and 15 ng/μL DNA. qPCR was run on a Quantstudio 5 (Thermo Fisher Scientific) with 2 minutes at 50°C, 10 minutes at 95°C then 40 cycles of 15 seconds at 95°C and 60 seconds at 60°C (strong stop, second strand transfer, GFP) or 10 minutes at 95°C then 50 cycles of 15 seconds at 95°C and 90 seconds at 60°C (2LTR circles). Analysis was carried out using Thermo Fisher Design and Analysis software.

To measure ISGs by qPCR, RNA was extracted from cells RNeasy purification kit (Qiagen, 74106) following the manufacturer’s instructions, including on-column DNase digest for 15 minutes at room temperature using RNase-Free DNase Set (Qiagen, 79254). To convert this to cDNA, 500 ng RNA in 11 μL was mixed with 1 μL 10 mM dNTP (Thermo Fisher Scientific, R0191) and 1 μL oligo(dT) primer (Thermo Fisher Scientific, SO132) and incubated at 65°C for 5 minutes and then transferred to ice for 1 minute. This was mixed with 4 μL 5x first-strand buffer (Invitrogen, Y02321), 1 μL 0.1M DTT (Invitrogen, Y00147), 1 μL RNase OUT (Invitrogen, 10777019) and 1 μL Superscript III reverse transcriptase (Invitrogen, 18080044) and incubated at 50 °C for 60 minutes then 70°C for 15 minutes.

qPCR was then performed on cDNA diluted 1 in 5 using Fast SYBR Green PCR master mix (Applied Biosystems, 4385612) and primers detailed below in a 384-well plate. qPCR was run on a Quantstudio 5 (Thermo Fisher Scientific) with 20 seconds at 95°C then 40 cycles of 1 second at 95°C and 30 seconds at 60°C. Analysis was carried out using Thermo Fisher Design and Analysis software. Gene expression was normalised to GAPDH control and DMSO treated samples.

#### Immunoblotting

Cells were lysed in RIPA buffer (50 mM Tris pH 8.0, 150 mM sodium chloride, 1.0% Triton X-100, 0.5% sodium deoxycholate, 0.1% SDS with phosphatase (PhosSTOP, Roche) and protease (cOmplete mini EDTA-free, Roche) inhibitors) by incubation for 10 minutes on ice. Cell lysates were centrifuged for 10 minutes at 13200 rpm at 4 °C and supernatants mixed with 4X loading buffer (200 mM Tris pH 6.8, 8% SDS, 0.4% bromophenol blue, 40% glycerol and 5% beta-mercaptoethanol). For viral supernatants, around 800 μL were centrifuged at 14000rpm, 4 °C for 1.5 hours. Supernatants were removed, and pellets resuspended in 1X loading buffer. Samples were incubated at 95°C for 10 minutes. Proteins were separated by SDS-PAGE with 10 or 12% SDS polyacrylamide gels with 5% stacking gels prepared in-house. Proteins were transferred onto nitrocellulose membranes using Trans-Blot Turbo RTA Transfer Kit (BioRad, 1704270), following the manufacturer’s protocol. Membranes were blocked for 1 hour in 5% milk PBS-T (1X PBS, 0.1% Tween-20). Primary antibodies (1:2000 Ms Anti-p24 183-H12-5C HIV Reagents Program, 1:1000 Rb Anti-VSV-G ab19257 Abcam, 1:10000 Ms Anti-β-actin ab6276 Abcam, 1:1000 Rb cGAS, Santacruz) were incubated for 1 hour at room temperature or overnight at 4 °C. The membranes were washed three times in PBS-T, the incubated with secondary antibodies (1:10000 IRDye® 680LT Gt Anti-mouse 926-68020, LI-COR Biosciences, 1:10000 IRDye® 800CW Gt Anti-mouse 926-32210, LI-COR Biosciences, 1:10000 IRDye® 680LT Gt Anti-rabbit 926-68021, LI-COR Biosciences, 1:10000 IRDye® 800CW Gt Anti-rabbit 926-32211, LI-COR Biosciences) for 1 hour at room temperature. After washing three times with PBS-T and twice with PBS, proteins were visualised at 700nm and 800nm using a LI-COR Odyssey CLx imaging system. Densitometry was performed using Empiria Studio Software (LI-COR Biosciences).

#### Statistics

Statistical analysis was performed using GraphPad Prism and Origin Pro as indicted in figure legends.

## Data Availability

Model coordinates and structure factors for crystal structures have been deposited in the Protein Data Bank, under accession numbers: PDB-9RPC for Apo-CA, PDB-9S6J for CA-CPSF6, PDB-9S6N for CA-NUP153, PDB-9S6O for CA-JW3-76, PDB-9S7M for CA-JW3-93, PDB-6S6V for CA-JW3-94, PDB-9S6W for CA-JW3-100 and PDB-9S6X for JW3-134.

## Author Contributions

Conceptualisation: GJT, DLS, NP, KLM, MLG

Investigation: KLM, MLG, BG, JW, LH, LGT, LSN, JW, ET, RS, SO, NP

Funding acquisition: GJT, DLS, TB, DJ

Methodology: KLM, MLG, BG, JW, LH, LGT, DA, NP

Project administration: DLS, GJT

Supervision: NP, DLS, GJT, LGT

Visualisation: KLM, MLG, NP

Writing (original draft): KLM, MLG, BG, NP, GJT

Writing (review and editing): KLM, MLG, NP, GJT, DLS

## Acknowledgments

We thank Gabriel Waksman and Carolyn Moores for Birkbeck lab access and Chloe Orkin (QMUL) for helpful comments on the manuscript. This work was funded by Wellcome Investigator Award (220863) and Wellcome Discovery Award (325794) to GJT, Wellcome Collaborative award (214344) to GJT, DS, DJ, The European Research Council under the European Union’s Seventh Framework Programme (FP7/2007-2013)/ERC (grant HIVInnate 339223) to GJT and DS, National Health and Medical Research Council Ideas Grant (GNT2013215) to DJ and TB and Australian Research Council Discovery Grant (DP240102772) to DJ and TB. Synchrotron Access was supported by the iNEXT-Discovery proposal PID: 12462. The majority of our synchrotron data were collected at the EMBL Hamburg beamlines P13 and P14 at the PETRA III storage ring (DESY, Hamburg, Germany). We would like to thank all involved EMBL personnel for the assistance in accessing and using the beamlines. We also acknowledge the European Synchrotron Radiation Facility (ESRF, Grenoble France), in particular the personnel of the beamline ID23-1 for providing access to collect the CA-NUP153 dataset. Additionally, we thank the Diamond Light Source (DLS, Oxfordshire, UK) for access to beamline I04, where preliminary crystal screening was carried out and proved valuable to the success of this project.

